# Beta cell dysfunction occurs independently of insulitis in type 1 diabetes pathogenesis

**DOI:** 10.1101/2024.12.29.630665

**Authors:** Mollie K. Huber, Adrienne E. Widener, Alexandra E. Cuaycal, Dylan Smurlick, Elizabeth A. Butterworth, Nataliya I. Lenchik, Jing Chen, Maria Beery, Helmut Hiller, Ellen Verney, Irina Kusmartseva, Marjan Slak Rupnik, Martha Campbell-Thompson, Ivan C. Gerling, Mark A. Atkinson, Clayton E. Mathews, Edward A. Phelps

## Abstract

The loss of insulin secretory function associated with type 1 diabetes (T1D) is attributed to the immune-mediated destruction of beta cells. Yet, at onset of T1D, patients often have a significant beta cell mass remaining while T cell infiltration of pancreatic islets is sporadic. Thus, we investigated the hypothesis that the remaining beta cells in T1D are largely dysfunctional using live human pancreas tissue slices prepared from organ donors with recently diagnosed T1D. Beta cells in slices from donors with T1D had significantly diminished Ca^2+^ mobilization and insulin secretion responses to glucose. Beta cell function was equally impaired in T cell-infiltrated and non-infiltrated islets. Fixed tissue staining and gene expression profiling of laser-capture microdissected islets revealed significant decreases of proteins and genes in the glucose stimulus secretion coupling pathway. From these data, we posit that functional defects occur in the remaining mass of beta cells during human T1D pathogenesis.

## Introduction

The presence of islet autoantibodies (AAb) and a progressive loss of beta cell function are characteristics of type 1 diabetes (T1D) progression. Disease kinetics vary between patients with some individuals being diagnosed at only a few years of age while others have detectable AAb for several years prior to diagnosis^1–4^. Organ-specific immune cell infiltration into pancreatic islets (insulitis, including T cells, B cells, and macrophages) is considered a hallmark of T1D^5^. While facets of the islet-immune interface resulting in beta cell loss have been established in animal models of T1D, the sequence of immunological events from the initiation of autoimmunity to clinical diagnosis of T1D impacting islet function remains unknown in humans^6^.

Clinical studies have established that beta cells exhibit a period of declining function that begins months to years prior to diagnosis^7–14^. Early asymptomatic T1D patients show decreased insulin secretion responses to intravenous glucose^8,15,16^, while insulin release to other secretagogues is less perturbed^11,17^. These data align with histological analyses of tissue from organ donors, autopsies, or biopsies that frequently reveal abundant surviving beta cells at T1D diagnosis^5^. Indeed, endogenous C-peptide secretion can temporarily improve during the honeymoon period with the initiation of insulin therapy, and incretins like semaglutide and liraglutide can restore insulin independence for months to years after diagnosis^18–21^. The observation that beta cell function is reduced years prior to diagnosis has not been coupled with the concurrent presence of insulitis or a reduction in beta cell mass. Insulitis is observed most commonly in tissues from donors with recently diagnosed T1D (≤18 months) and rarely, in those positive for multiple AAb prior to T1D onset^5,22–25^. Furthermore, in cases where insulitis is found, it is relatively sparse, generally impacting a subset of insulin-positive islets^5,22,26^. These observations suggest that the clinical manifestations of T1D are the culmination of immune-mediated beta cell loss along with a parallel metabolic defect within the beta cells.

Here, we applied the live pancreas tissue slice model to investigate the *in-situ* functional and molecular changes that occur within human beta cells in T1D^27–30^. We studied organ donor tissue provided by the Network for Pancreatic Organ donors with Diabetes (nPOD) from individuals with no diabetes and AAb negative (ND: n=14), no diabetes and AAb positive (AAb+) with T1D-risk human leukocyte antigen (HLA) (n=11), and short-duration T1D (<4 years since diagnosis, n=9). Using dynamic Ca^2+^ imaging and insulin secretion assays, coupled with live staining for the beta cell surface marker, anti-ectonucleoside triphosphate diphosphohydrolase 3 (ENTPD3), and the T cell marker, CD3, we found that beta cell-containing islets in T1D donors universally failed to respond to glucose, despite retaining high viability and responsiveness to KCl. Importantly, islet dysfunction was not directly correlated with the presence of T cells, indicating that beta cells were dysfunctional regardless of local T cell infiltration on a per-islet basis. These findings suggest a widespread beta cell dysfunction that occurs in parallel with beta cell loss. A targeted assessment of the glucose-stimulated insulin secretion (GSIS) pathway identified differentially expressed genes and proteins involved in glucose import, glycolysis, oxidative phosphorylation, and membrane depolarization that correlate with beta cell dysfunction in T1D+ donors. Thus, we propose that metabolic dysfunction of remaining beta cells is a hallmark of T1D that contributes significantly to clinical symptoms at diagnosis.

## Methods

### Sample Collection

Live human pancreas tissue slices were received from the Network for Pancreatic Organ donors with Diabetes (nPOD) from donors classified by nPOD’s clinical team as ND (n=14), AAb+ (n=11), or confirmed cases of short duration (within 4 years of diagnosis) T1D (T1D+, n=9). Formalin-fixed, paraffin-embedded (FFPE) pancreas tissue sections used for staining were received from nPOD from ND donors (n=9), donors with a single AAb without T1D (sAAb+, n=7), donors with multiple AAb without T1D (mAAb+, n=7), and donors with confirmed cases of short duration (within 4 years of diagnosis) T1D (T1D+, n=6). Ten slides were received per donor. Optimal cutting temperature (OCT) pancreas tissue slides used for laser-capture microdissection were received from nPOD from ND (n=10), sAAb+ (n=3), mAAb+ (n=3), and donors with confirmed cases of T1D (within 7 years of diagnosis, T1D+, n=6). Donor characteristics and corresponding experiments are listed in **Supplementary Table 1**.

### Live Pancreas Tissue Slice Studies

Live human pancreas tissue slices were cultured overnight in slice media (HyClone Dulbecco’s Low Glucose modified Eagles Medium +L-Glutamine +Pyruvate, Fisher Scientific SH30021.01, 10% fetal bovine serum, heat inactivated Sigma F4135, 25 kIU/mL aprotinin from bovine lung BioUltra, 3-8 TIU/mg solid, >98%, Sigma A6106-100MG, and 1% antibiotic-antimycotic solution, Corning, 30-004-CI) at 24°C. Prior to staining and functional studies, slices were transferred to an incubator set to 37°C for at least 1 hour. After warming, slices were washed once in Krebs-Ringer bicarbonate HEPES buffer (KRBH) with 3 mM glucose (3G, low glucose). Slices were then stained individually in 8-well Ibidi dishes with anti-ENTPD3 (R&D Systems Human CD39L3/ENTPD3 Antibody AF4400) to identify beta cells, Ca^2+^ indicator (Fluo-4, AM, Thermo Fisher Scientific F14201 or Calbryte 520 AM, AAT Bioquest 20650) to detect changes in intracellular Ca^2+^, and anti-CD3 (Biolegend Alexa Fluor 647 anti-human CD3 300416 Clone UCHT1) to identify and track endogenous T cells. Viability was assessed via SYTOX Blue (Thermo Fisher Scientific 501137613) staining. Live imaging studies were conducted using a Leica SP8 confocal microscope equipped with a live cell incubation system allowing for long-term stable recordings with continual perfusion via programmable syringe pumps. Slices were initially perfused in KRBH buffer with 3G, then stimulated with 16.7 mM glucose (16.7G, high glucose), and finally, with 30 mM potassium chloride (KCl). The timing of each stimulation protocol varied slightly between individual cases and can be found in **Supplementary Table 2**.

### Ca^2+^ Recording Analysis

Ca^2+^ recordings of ENTPD3+ islets within slices were motion and bleach corrected using the StackReg plugin for the FIJI distribution of ImageJ^31,32^. Regions of interest (ROIs) were hand-selected over individual beta cells using ENTPD3 positivity or reflected light for six early cases. Ca^2+^ indicator mean fluorescence intensities (MFIs) for the cells represented within ROIs were calculated and exported from ImageJ^33^. The Ca^2+^ indicator fluorescence of each beta cell was normalized to the mean of the first 10-100 frames of the recording (F/F_0_), depending on the frame rate. Traces and dynamics were then visualized and analyzed in RStudio v2024.04.2^34^.The peak Ca^2+^ response of each recording was determined by averaging all beta cell traces and finding the maximum MFI in the 3G, 16.7G, and KCl stimulation periods. The high glucose over low glucose ratio (HG/LG) was calculated by dividing the average MFI in the 16.7G region by the average MFI of the 3G region for each beta cell trace in the islet. From the HG/LG calculation, a ratio of 1.2 was set as the threshold of a functional response. The number of beta cell that met or exceeded a HG/LG ratio of 1.2 was divided by the total number of beta cells to determine the fraction of responding beta cells. Peak Ca^2+^ response, HG/LG ratio, and fraction of responding cells were analyzed by fitting a linear mixed effects model with a restricted maximum likelihood (REML) method and considering the case number as the random effect using the nlme package in R^35,36^. Linear mixed effects models were then analyzed using a one-way ANOVA and Tukey’s post hoc testing using the lsmeans package in R^37^. Significance was determined as a 95% confidence interval and p < 0.05. This statistical analysis approach was used to take into account the hierarchical nature of the dataset with multiple beta cells per islet, multiple islet recordings per donor, and multiple donors per donor type in accordance with suggested best practices for communicating reproducibility and variability in cell biology^38^. One recording from nPOD case 6590 (ND) was excluded from analysis due to erratic Ca^2+^ flux patterns that were independent of stimulations, heatmap can be found in **Supplementary Figure 4**. Ca^2+^ analysis code is available at https://github.com/PhelpsLabUF/Slice_Calcium_Imaging_Visualization_Analysis.

### Slice Perifusion

Perifusion studies were conducted using a Biorep Technologies Perifusion System (Biorep Technologies, Cat. No. PERI-4.2). Live human pancreas tissue slices were rested at room temperature on a plate rocker set to 30 RPM for a minimum of 1.5 hours in 3G with aprotinin (1:100). Three slices each were placed into three closed perifusion chambers and connected to the system. Tissue slices were perifused at a flow rate of 100 μL/min at 37°C with KRBH supplemented with 3G, 16.7G, 1 mM glucose (1G), then 60 mM KCl, and the perifusate was collected in 96-well plates at 1-minute intervals. Perifusates were stored at −20°C until commercial insulin ELISAs were run^28,39^. Insulin measurements from perifusion experiments were analyzed with a custom R script. Briefly, stimulation times for each stimulation were corrected with a three-minute delay empirically determined by the nPOD team. Data were normalized to the average insulin measurement from the low-glucose (3G) stimulation, and area under the curve (AUC) was calculated for each stimulation condition. AUC measurements were divided by the stimulation time to obtain AUC/min per stimulation condition.

### T Cell Quantification

T cells were quantified manually from the three-dimensional Z-stacks of live islets captured in Ca^2+^ recordings. T cells were identified through positive staining from an anti-CD3 antibody (Biolegend Alexa Fluor 647 anti-human CD3 300416 Clone UCHT1) and were counted if they were within one cell body length of the islet boundary or residing within a focal or infiltration insulitis lesion. T cell tracking was performed using the manual tracking mode for the TrackMate plugin for ImageJ^40^.

### Immunofluorescent Staining of Fixed Pancreas Tissue Sections

FFPE sections (4 µm) were adhered on Superfrost Plus slides (Fisher Scientific 12-550-15) and stored at room temperature until use. For deparaffinization, slides were heated for 30 minutes at 65°C followed by submersion in Histoclear II (Electron Microscopy Sciences 64111-04) twice for five minutes. Sections were then rehydrated using 100% ethanol (twice for two minutes), 95% ethanol (dipping four times, then soaking for three minutes), 70% ethanol (one minute), and deionized water (one minute).

Antigen retrieval (AR) was performed in a steamer as follows: coplin jars were filled with AR buffer (Borg Decloaker, BioCare BD1000G1) and placed in a steamer (Black and Decker, HS1050) for 15 minutes to preheat. Slides were then placed in the heated AR solution and steamed for 20 minutes. The coplin jars were removed from the steamer and placed on the counter to cool for 20 minutes. After cooling, slides were washed in deionized water for two minutes. A hydrophobic barrier pen (Vector Laboratories H-4000) was used to draw borders around the tissue section being careful not to let the section dry out. Sections were then washed in TBS with Tween (TBST, Thermo Scientific, J77500.K8) twice for five minutes each followed by incubation in blocking buffer (Opal Antibody Diluent/Block, Akoya Biosciences ARD1001EA) for ten minutes.

In order to stain for all target proteins, antibodies were split into 10 four-marker panels, with CD3 and insulin present in all panels. Primary and secondary antibodies used and their corresponding protocols are listed in **Supplementary Table 3.** Both primary and secondary antibodies were diluted in Opal Antibody Diluent/Block and a minimum of 200 µl antibody solution was used per section for staining. Antibodies were stained sequentially. Generally, the first primary antibody was added to the section after blocking, and slides were incubated in a humidified chamber for 16 hours at 4°C followed by one hour at room temperature. Sections were washed using a gentle stream of TBST buffer delivered using a squirt bottle to remove the majority of the primary antibody solution and placed in a coplin jar filled with TBST to wash on a nutating mixer (Fisher Scientific 88-861-041) for 30 minutes. The secondary antibody solution was added and sections were incubated in a black slide humidity chamber (FisherScientific NC9062083) for ten minutes. Sections were washed three times with a gentle stream of TBST for five minutes each. Sections were incubated in blocking buffer for 10 minutes to prepare for the next primary antibody in the sequence, and the incubation and washing process was repeated for the second and third primary and secondary antibody combinations. On the final day of staining after the third secondary antibody was added, slides were again washed with TBST. The conjugated insulin antibody (BD Pharmingen 565689) was added and the slides were incubated at room temperature for one hour. Nuclei were counterstained using Hoechst S769121 (Nuclear yellow, 1µM stock solution prepared with deionized water (ab138903) and aliquots stored at −20°C) diluted (1:50) for ten minutes. Slides were washed, and coverslips were mounted with ProLong Gold Antifade Mountant (Invitrogen P36930). Slides were left to dry overnight in the dark at room temperature and stored in opaque slide boxes at 4°C prior to imaging.

Slides were imaged using a Leica SP8 confocal microscope and a Leica Stellaris 5 confocal microscope. The microscope and imaging settings were kept consistent within each panel. 30 islets were imaged per slide with the exception of some slides from T1D+ donors where fewer than 30 insulin-positive islets were present in the tissue section. For these cases, all islets present were imaged.

### Confocal Image Quantification

Islet images were analyzed using CellProfiler^41^. Insulin positive cells were identified as primary objects. The area was then masked to identify the islet area for MFI measurements. The beta cell area was expanded to create a secondary object and inverted to mask the exocrine area for MFI measurements. MFIs were measured for both the insulin-positive islet area and the exocrine area for the different markers. MFI values were scaled from 0-1 by CellProfiler and exported into Excel. The averages for individual islets and cases were calculated for each marker within both the islet and exocrine areas. The islet averages were analyzed by fitting a linear mixed effects model by an REML method^35,36^. The models were then analyzed using a one-way ANOVA and Tukey’s post hoc testing, with a significance of p < 0.05^37^. MFI values were also visualized collectively using a heatmap. Heatmap visualization of scaled (0-1) intensity data was performed in RStudio v2024.04.0 with the ComplexHeatmap package v2.20.0^42,43^. Scaled intensity data was z-score transformed considering the ND group as the baseline population for mean and standard deviation (SD) calculations. CD3+ and CD45+ cells were both counted manually and were only counted if the cells were infiltrating the insulin-positive area of the islet.

### Microarray Gene Expression Analysis

Fresh frozen pancreas sections were obtained from ND (n=10), sAAb+ (n=3), mAAb+ (n=3), and T1D+ donors (n=6). Blocks were selected based on immunohistochemistry staining and contained insulin positive islets and/or T cell infiltration as defined by six or more CD3+ cells immediately touching the islet endocrine cells^5^. Blocks were cut into thick (10 μm) serial (3-4) sections and placed onto PEN-slides (Leica) along with thin (4 μm) sections placed onto Superfrost slides (Thermofisher) before and after the thick sections. Thin pancreas sections were immunostained for insulin and CD3 to identify islets with residual beta cells and T cell infiltration. Thick sections were used to laser-microdissect single islets from dehydrated slides. Laser microdissections yielded 3-4 sections per islet captured into 0.6 mL RNAse-free tubes. Sets of 7-25 islets were laser-captured per donor. Total islet RNA was extracted using the PicoPure RNA kit (ThermoFisher). DNAse treatment was performed using recombinant DNase I (RNase-free DNAse kit, Worthington Biochemical Corporation). A 4 μL aliquot was used for RNA quality control (QC) assays: the RNA concentration and quality (RIN) were assessed using an Agilent 2100 Bioanalyzer (Agilent Technologies) at the University of Florida Interdisciplinary Center for Biotechnology Research (ICBR). The remaining sample (8-12 μL) was stored at −80°C until batch shipment on dry ice to the University of Tennessee Health Science Center, Memphis, TN where microarray gene expression profiling was performed with the Affymetrix Human Gene 2.0 ST expression array (Thermo Fisher Scientific). Gene expression analysis was performed in RStudio v2024.04.0. The SST-RMA-normalized microarray signals were variance-stabilized using the justvsn function implemented in the vsn package v3.72.0^44^ before statistical testing.

QC checks were performed with Uniform Manifold Approximation and Projection (UMAP) algorithm (M3C package v1.26.0), demonstrating overall good clustering of islets across the four clinical phenotypes (**Supplementary Figure 1**)^45^. Initial differential expression (DE) analysis with INS+CD3-islets revealed the expected presence of non-beta cell transcripts. Hence, islets enriched for three or more alpha-cell specific transcripts (z-score > 1) were removed from the dataset. In addition, only islets with residual INS+ beta cells were used for subsequent analyses. Gene expression analyses presented here were performed with INS+CD3-islets only. The final dataset consisted of 34 ND islets, 13 sAAb+ islets, 18 mAAb+ islets, and 13 T1D+ islets.

Individual islet gene expression for each marker of interest was obtained and averaged for each donor across the four clinical phenotypes. Donor average gene expression values were plotted using GraphPad Prism software version 10.3.1. Statistical significance was defined as p < 0.05 after a one-way ANOVA with Tukey’s multiple comparisons test. Gene expression data are available at Gene Expression Omnibus (GSE284772).

### Ethical Approvals

All procedures using human slices were performed according to the established standard operating procedures of the nPOD/OPPC and approved by the University of Florida Institutional Review Board (IRB201600029) and the United Network for Organ Sharing (UNOS) according to federal guidelines, with informed consent obtained from each donor’s legal representative. Pancreas tissue from human donors with or without T1D was procured from the nPOD program at the University of Florida (RRID: SCR_014641, https://www.jdrfnpod.org). For each donor, a medical chart review was performed, and C-peptide was measured, with T1D diagnosed according to the guidelines established by the American Diabetes Association (ADA). Demographic data, hospitalization duration, and organ transport time were obtained from hospital records. Donor pancreata were recovered, placed in transport media on ice, and shipped via organ courier to the University of Florida. The tissue was processed by a licensed pathology assistant. Detailed donor information is listed in Supplementary Table 1. nPOD tissues specifically used for this project were approved as nonhuman by the University of Florida IRB (NH00041892, NH00042022).

## Results

### ENTPD3 Staining Identifies Beta Cells in Live Pancreas Tissue Slices

A period of beta cell dysfunction during T1D development has been well-established in clinical studies ^7,8,10–14^. However, this dysfunction has been understudied in isolated islets; thus, the mechanism(s) driving it has remained unclear. Islets are difficult to isolate from T1D+ donors due to the islet degradation that is associated with disease^46^. Additionally, the isolation procedure results in islets being removed from their microenvironment causing them to lose their underlying pathologies and associated immune cells. In the small number of studies performed on isolated human T1D+ islets, beta cells have been found to persist in T1D+ donors despite diminished responses to high glucose ^47–49^. Another study investigating T1D+ islets found that high glucose responses were maintained when normalized to insulin content with fewer beta cells being present within the islets ^50^. However, this persisting function can likely be attributed to the best and most intact islets being most likely to survive the isolation procedure. Live human pancreas tissue slices present a unique opportunity to study islets at the different stages of T1D development including islets with high levels of immune cell infiltration and structural degradation. This combined with the preservation of the native microenvironment (**Figure 1a**) makes the model optimal for investigating beta cell function in T1D ^27–30,33,39,51–58^.

**Figure 1:**
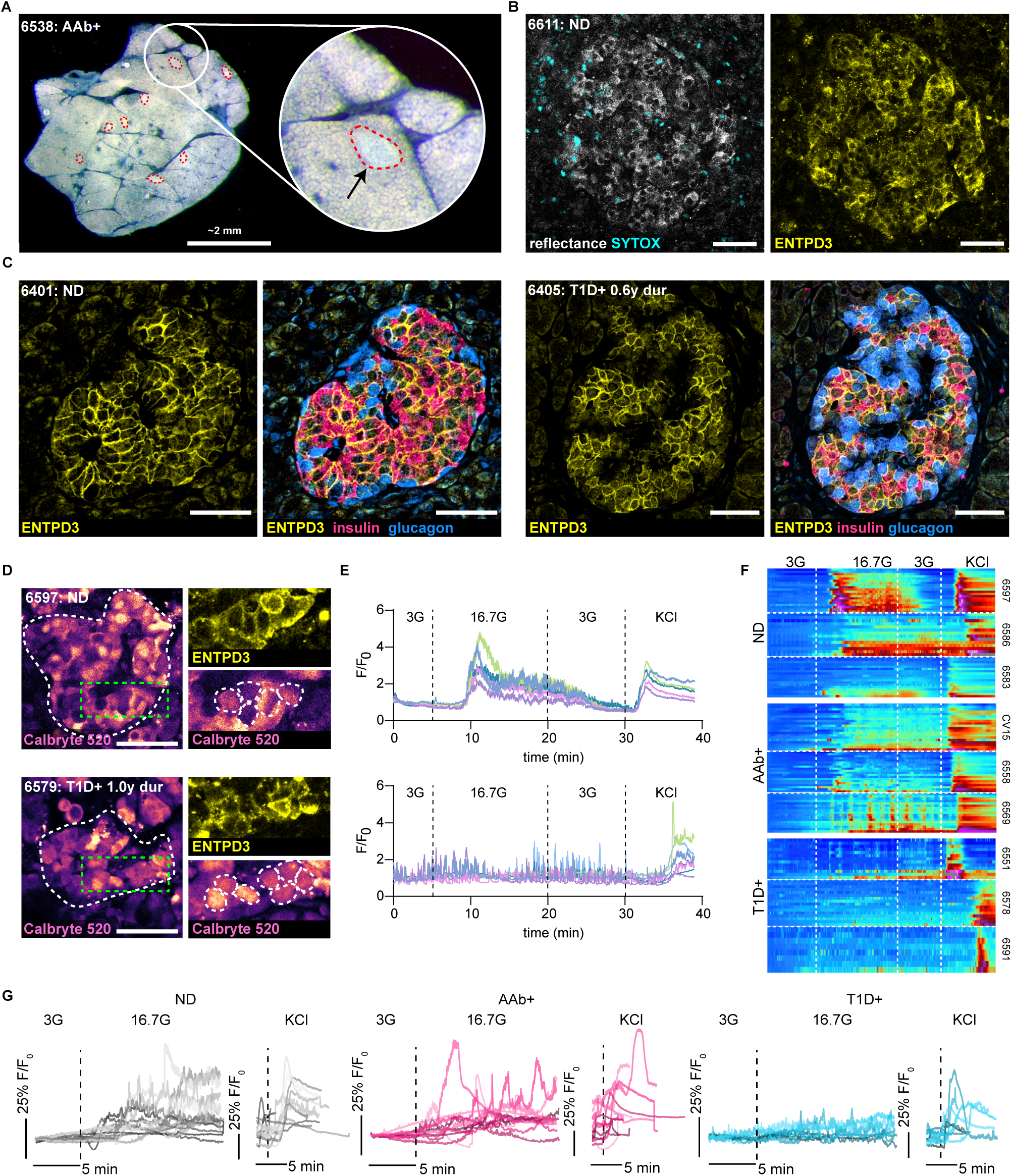
ENTPD3 staining identifies beta cells in live pancreas tissue slices from ND and T1D+ donors. (**A**) Darkfield stereomicroscopy image of a live human pancreas tissue slice with islets outlined in red. (**B**) Confocal image of an islet within a live slice indicated by reflected light and ENTPD3. Dead cells are indicated by SYTOX staining. (**C**) Fixed tissue staining of FFPE human pancreas tissue sections showing ENTPD3 positive cells co-localize with insulin positive cells and not glucagon positive cells in both ND and T1D+ cases. (**D**) Reflective and ENTPD3 positive islets within ND and T1D+ slices outlined in red. Still frames of Ca^2+^ responses to high glucose (16.7G) of the same islets outlined in white (middle). Magnified regions outlined in green within these islets showing individual ENTPD3+ cells, outlined in white. (**E**) Heatmaps depicting representative islet Ca^2+^ responses to low glucose (3 mM), high glucose (16.7 mM), and KCl (30 mM) in three donors from ND, Aab+, and T1D+ cases. Each column represents the Ca^2+^ response in an individual ENTPD3+ cell. (**F**) Traces of the mean islet Ca^2+^ response from all recorded ND (n=14), Aab+ (n=11), and T1D+ cases (n=9). Scale bars, 50 µm.

To investigate beta cell function in T1D+ slices it is necessary to identify islets that still contain viable beta cells. A beta cell surface marker is needed as intracellular markers, such as insulin, cannot be stained in live tissue. NTPDase3 (ENTPD3) has been identified as a reliable beta cell surface marker in flow cytometry, fixed tissue sections, and whole islets ^59,60^. Here, we conjugated a fluorophore to an anti-ENTPD3 antibody to label beta cells within the slices and found effective cell surface labeling of live islets in slices from ND donors (**Figure 1b**). Fixed tissue staining revealed strong co-localization between ENTPD3 and insulin along with a lack of colocalization between ENTPD3 and glucagon (**Figure 1c**). While ENTPD3 has been shown to also colocalize in some delta cells^59^, here we clearly demonstrate that it can be successfully used to identify beta cells within slices. ND ROIs gated on individual ENTPD3^+^ cells exhibited typical beta cell glucose and KCl cytosolic Ca^2+^ dynamics (**Figure 1d-e**). Taken together with the existing literature on ENTPD3 surface expression on beta cells, we conclude that ENTPD3 is a reliable surface marker to identify beta cells in pancreas slices without perturbing their function.

Next, we tested whether ENTPD3 can be used to identify remaining beta cells in slices from donors with short-duration T1D. We identified numerous ENTPD3^+^ cells within islets from T1D+ donors, though, as expected, less frequently than in ND donors. Fixed tissue staining showed ENTPD3 colocalization with insulin and a lack of colocalization with glucagon was preserved in T1D+ donors (**Figure 1c**), indicating that the labeled cells are indeed beta cells. The ENTPD3^+^ cells in T1D+ slices typically failed to respond to high glucose yet had preserved responses to KCl (**Figure 1d-e**). As this result has not previously been subject to extensive investigation, we were motivated to study beta cell function in a larger cohort of slice donors, including additional T1D+, sAAb+, and mAAb+ donors to understand how beta cell function relates to the presence of infiltrating CD3^+^ T cells.

Beta cell Ca^2+^ responses to 16.7G and 30mM KCl stimulation in ND slices are represented in individual beta cells plotted as heatmaps (**Figure 1f**) and in average traces of all beta cells within an islet with trace colors representing individual cases (**Figure 1g**). These beta cell responses were maintained in AAb+ slices at both the cellular (**Figure 1f**) and islet level (**Figure 1g**). However, we found that high glucose responses were lost within T1D+ slices when looking at individual beta cell behavior within an islet (**Figure 1f**) and in islets overall (**Figure 1g**). KCl responses were maintained in these T1D+ cases, emphasizing that some beta cells are alive but do not respond appropriately to high glucose stimulation.

### Beta Cells Exhibit a Period of Dysfunction Prior to Death

To further characterize the beta cell responses within our cohort, we analyzed the Ca^2+^ recordings in several ways. To assess if there were any differences in activity between groups, we compared the peak Ca^2+^ response during low glucose, high glucose, and KCl stimulations. There were no significant differences in low glucose (3G) basal activity between donor types (**Figure 2a**). However, there was a significantly diminished response to high glucose (16.7G) in T1D+ beta cells when compared to both ND and Aab+ donors (**Figure 2b**). There were no significant differences between peak Ca^2+^ responses to KCl (**Figure 2c**), indicating dysfunction in the glucose metabolism pathway in remaining beta cells from T1D+ donors. These dysfunctional responses to high glucose are further emphasized when we took the ratio of high glucose to low glucose responses, allowing better visualization of each beta cells’ ability to respond to high glucose stimulation over baseline. When the ratios were compared between the different donor groups, T1D+ cases again had significantly diminished responses, indicating their general inability to respond above baseline to glucose stimulation with individual beta cells shown in **Figure 2d**. Finally, we assessed the fraction of the total beta cells within each islet that responded to high glucose stimulation. Most beta cells within ND and AAb+ slices were responsive to high glucose, while only a minority of islets in T1D+ slices contained any high glucose-responsive beta cells, with most T1D+ islets having no beta cells responsive to high glucose (**Figure 2e**). We also confirmed that the observed functional defect was not driven by sex differences between donor groups (**Supplementary Figure 2**). The trend of beta cell responses in the different donor types is further demonstrated by heatmaps with superimposed average traces from all Ca^2+^ recordings within each donor group (**Supplementary Figures 3-5**). While the strength and length of the responses vary, both ND (**Supplementary Figure 3**) and AAb+ (**Supplementary Figure 4**) slices generally maintained prominent responses to both high glucose and KCl stimulations. Conversely, high glucose responses were generally lost in T1D+ slice recordings, while KCl responses remained strong (**Supplementary Figure 5**).

**Figure 2:**
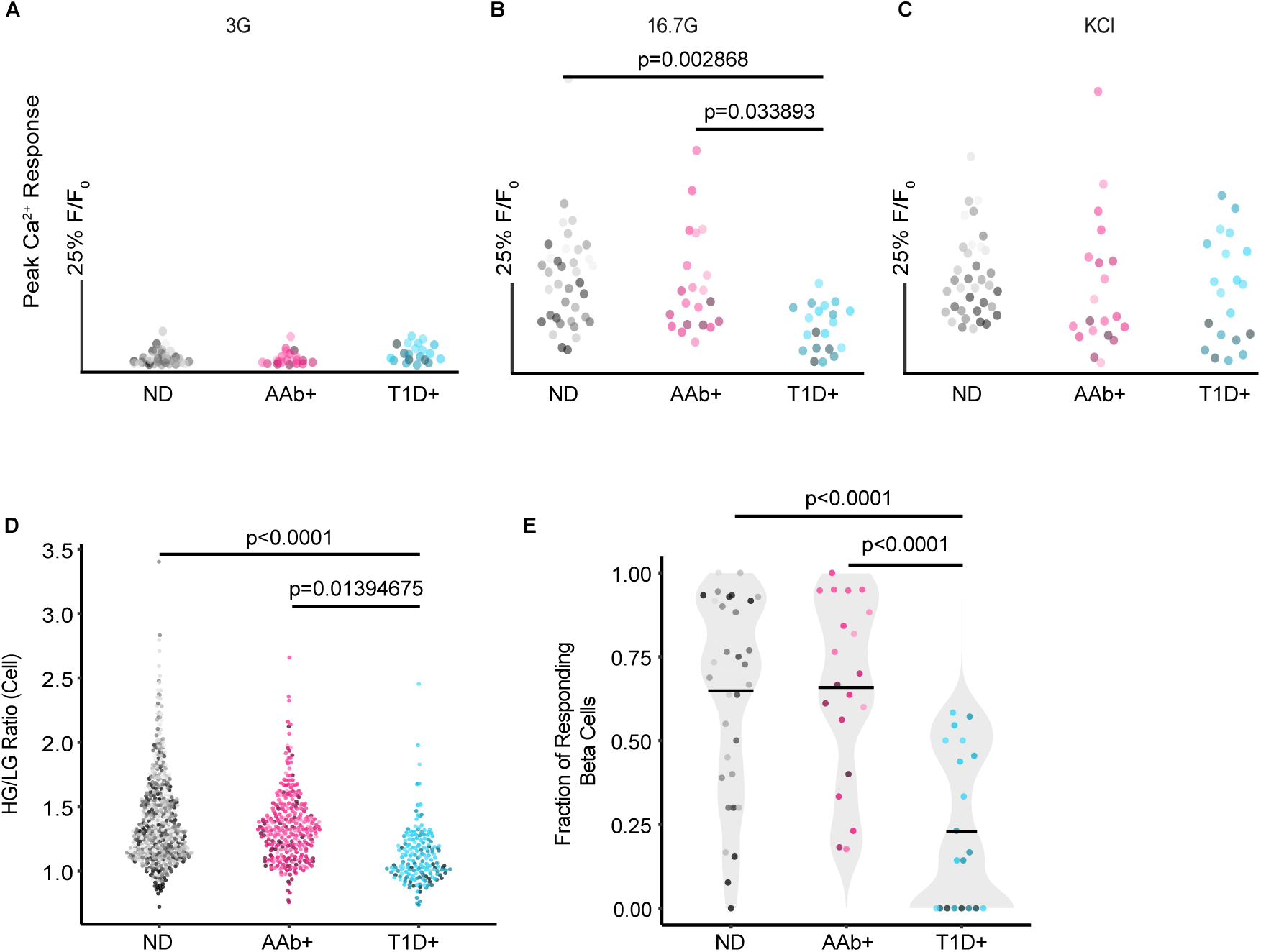
Beta cells exhibit a period of dysfunction prior to death. (**A-C**) The maximum Ca^2+^ response of the mean islet trace during basal activity in low glucose (3 mM), high glucose (16.7 mM), and KCl (30 mM). Each dot represents an islet recording and the shade represents donor for ND (n=14), Aab+ (n=11), and T1D+ (n=9) slice cases. (**D**) The ratio of the high glucose response to the low glucose response for individual beta cells in ND, Aab+, and T1D+ Ca^2+^ recordings demonstrates a loss of high glucose responses in T1D+ donors. Each dot represents a beta cell. (**E**) The fraction of beta cells responding to high glucose within each islet recorded from ND, Aab+, and T1D+ cases. Each dot represents an islet recording. Center line indicates the mean. One-way ANOVA with multiple comparisons.

We also monitored insulin secretion by dynamic perifusion of slices in low glucose, high glucose, and KCl and analyzed the fractionated effluent by ELISA for insulin (**Figure 3**). The loss of glucose-responsive Ca^2+^ signaling in T1D was mirrored by a similar loss of both first- and second-phase insulin secretion despite preservation of an insulin secretion response to KCl, verifying that these slices from T1D+ donors, with relatively short disease duration, retained a reserve of viable insulin-producing beta cells. These data establish a consistent trend of dysfunctional glucose responses in remaining beta cells in T1D+ slices. We next wanted to investigate whether T cell infiltration drives this dysfunction, as this is a critical component of understanding T1D pathogenesis.

**Figure 3:**
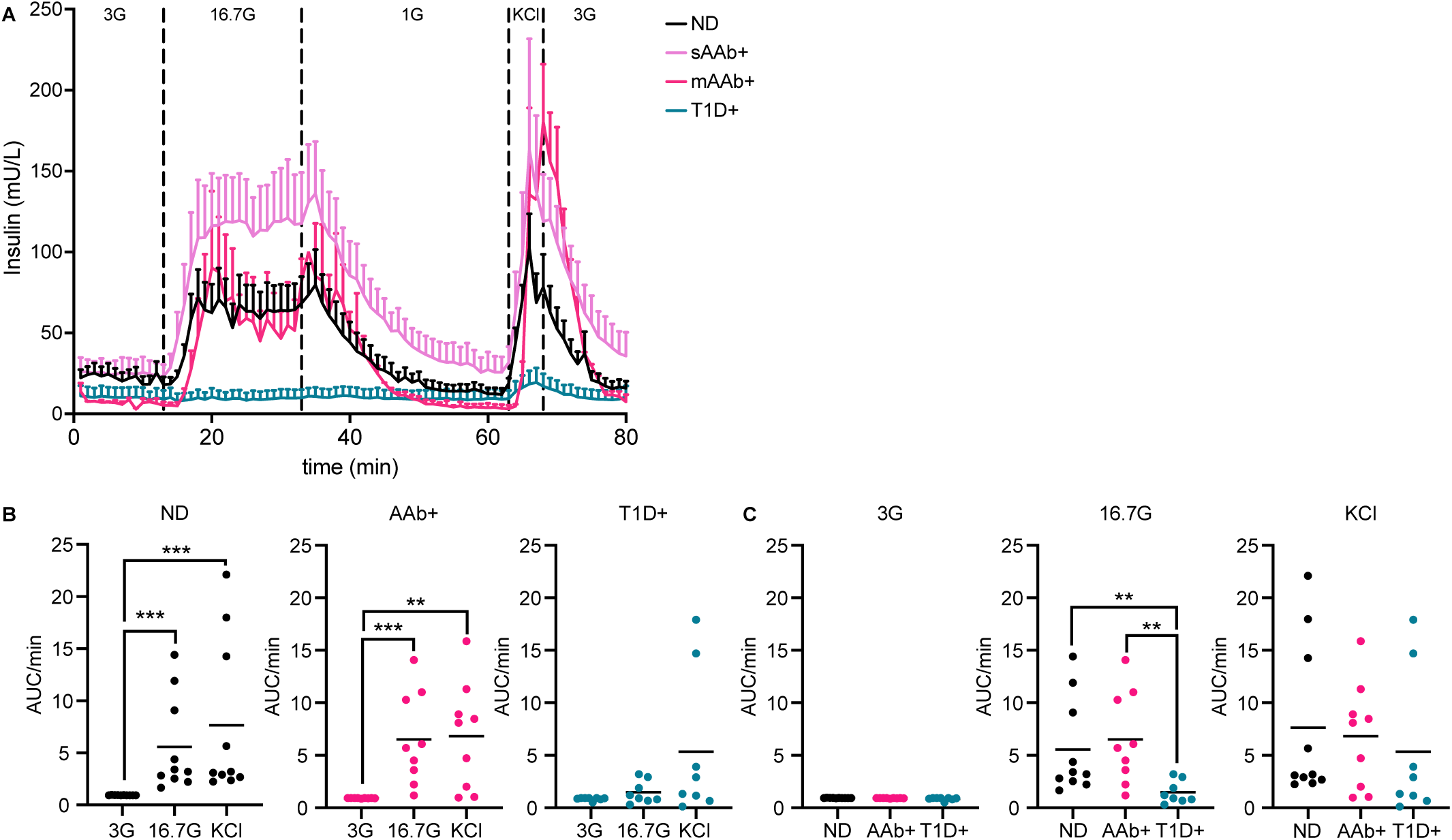
Beta cell insulin secretion decreases after T1D diagnosis. (**A**) Average insulin secretion traces from ND, Aab+, and T1D+ donors to low glucose (3 mM), high glucose (16.7 mM) and KCl (30 mM). (**B**) Quantification of AUC per minute of insulin secretion of live pancreas tissue slices from ND (n=10), Aab+ (n=9), and T1D+ (n=8) donors. Each dot represents one donor. (**C**) Quantification of AUC per minute of insulin secretion of the donor groups during different glucose stimulations. One-way ANOVA with multiple comparisons run on log10 transformed data, **P<0.01, ***P<0.001.

### Beta Cells in T1D are Dysfunctional Regardless of T Cell Infiltration

T cell infiltration into islets and the resultant destruction of beta cells are well-established components of T1D pathogenesis^5,22,23,25,61^. However, the studies investigating insulitis in human tissues have been predominantly conducted in fixed samples. As a result, the impact of endogenous T cell infiltration on beta cell function has yet to be determined. The live pancreas tissue slice model is ideal for addressing this gap as the beta cells remain viable and functional with preserved underlying tissue pathologies^27–30,55,62^. As such, we combined our beta cell functional assessments with simultaneous T cell recordings. Within our T1D+ donor cohort, insulitic,(at least 15 infiltrating CD45+ cells per islet in at least 3 islets along with the presence of insulin negative islets) insulin+ islets were identified in digital histology slides from the nPOD histopathology database https://portal.jdrfnpod.org/. We identified ENTDP3+ islets in our T1D+ donor cohort that were highly CD3+ T cell infiltrated through immunofluorescent staining of the live slices (**Figure 4a**). We captured several timelapse recordings of live endogenous CD3+ T cells interacting with ENTPD3+ islets in the first known live recordings of human insulitis *in situ* (**Supplementary Videos 1-3**).

**Figure 4:**
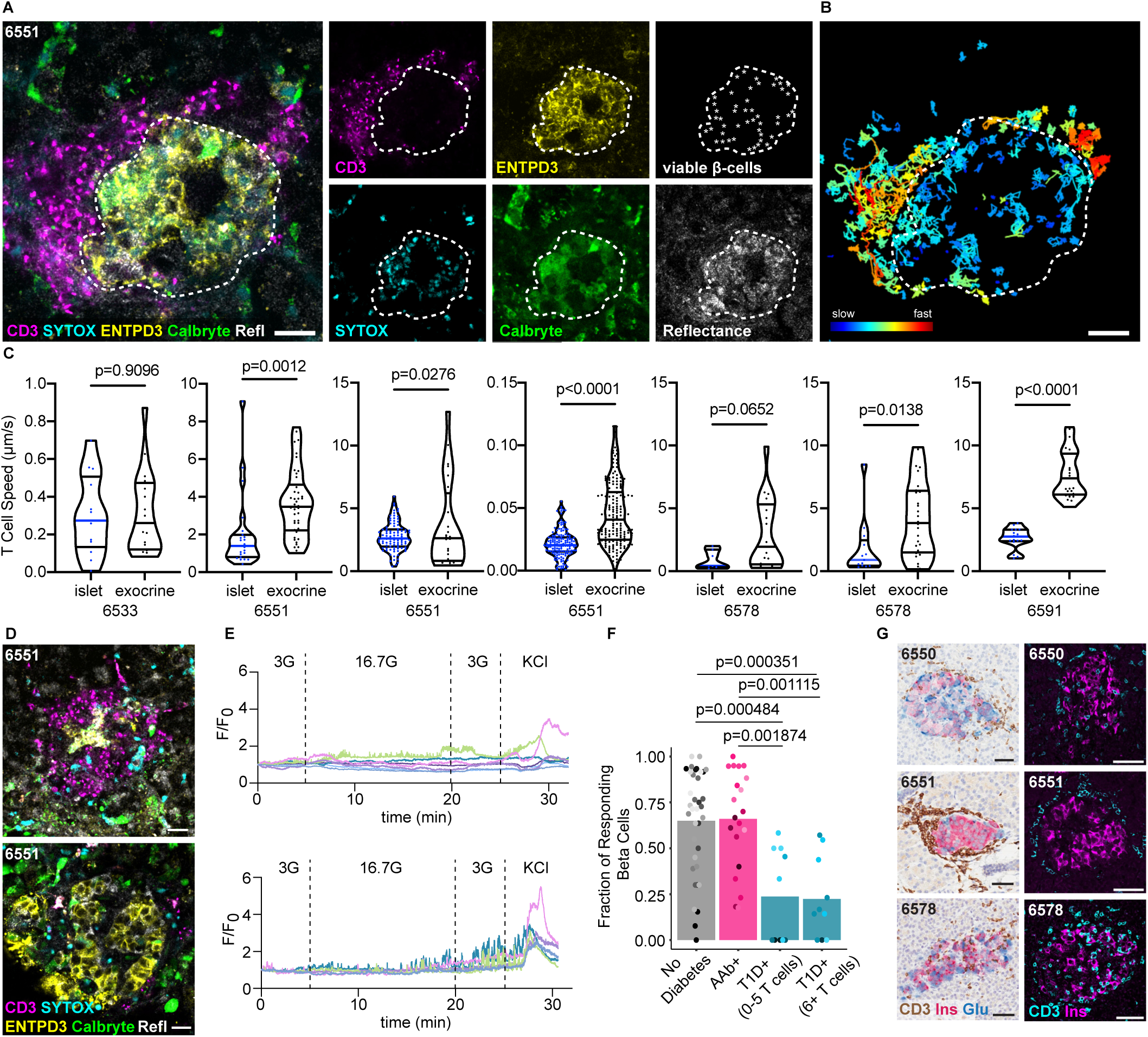
Beta cells in T1D are dysfunctional regardless of T cell infiltration. (**A**) Still image from a confocal timelapse recording of live endogenous insulitis within a T1D+ human pancreas tissue slice. Individual channels indicate the presence of viable beta cells (ENTPD3+/SYTOXneg), and CD3+ T cells. Scale bars, 20 µm. (**B**) Cell tracks demonstrating CD3+ T cell movement within and around the insulitic islet over 30 minutes. Scale bars, 20 µm. (**C**) Quantification of T cell motility within several different T cell infiltrated T1D+ islets compared to exocrine tissue. Unpaired t test. (**D**) Confocal images of CD3-infiltrated and non-infiltrated islets from the same T1D+ donor. Scale bars, 20 µm. (**E**) Ca^2+^ traces of viable ENTPD3+ cells in the depicted infiltrated (top panel) and non-infiltrated (bottom panel) T1D islets. (**F**) The fraction of beta cells responding to high glucose (16.7G) within each islet recorded for ND, Aab+, and T1D+ slices. T1D cases are separated into islets with low or high numbers of T cell infiltrates. ND and Aab+ islets all have five or fewer T cells. Stats to be added/ updated after analysis (**G**) Images of FFPE histological sections from T1D+ donors confirming the presence of insulin+ islets with insulitis in the studied donors. Scale bars, 50 µm.

Timelapse recordings of CD3+ infiltrated islets captured in four of the T1D+ donors in our cohort showed high levels of T cell motility within and around ENTPD3+ islets. The amount of infiltration was heterogeneous both within the same donor and between donors ranging from a high number of T cells (40+) swarming around and within the islet to more localized infiltration of 6-10 CD3+ T cells within the islet and at the islet periphery. We observed instances of T cells physically manipulating and disrupting beta cells which may constitute individual beta cell killing, although complete islet destruction did not occur within the time periods of our live observations (30 minutes to 2 hours). We performed cell tracking analysis and observed marked differences in T cell migration speed where T cells inside the islet border migrated more slowly than T cells outside the islet periphery (**Figure 4b**). A similar difference in cell migration speed was observed in all heavily infiltrated islets (7 islets in 4 T1D donors) (**Figure 4c**).

Studies conducted by Postić et al. established the presence of beta cell dysfunction prior to heavy immune cell infiltration in slices generated from NOD mice^58^. As we have established that human beta cells in T1D+ donors are alive but have dysfunctional responses to high glucose stimulations, we next investigated whether T cell infiltration was correlated with this dysfunction. Unlike the NOD mouse, a minority of islets within the human pancreas have insulitis at T1D onset^5^. Thus, CD3+ infiltrated (>6 T cells), ENTPD3+ islets were present but rare in slices from T1D donors, which allowed their functional comparison with uninfiltrated islets in the same donor (**Figure 4d**). We quantified the number of islet-infiltrating CD3+ T cells present in each Ca^2+^ recording and plotted those values against the fraction of beta cells responsive to high glucose for each donor group (**Figure 4f**). We found that the islets were equally dysfunctional regardless of whether they contained few (0-5) or many (6+) infiltrating T cells (**Figure 4f**), even within islets from the same donor (**Figure 4d,e**). We confirmed that islets in the slices recorded from these donors and in fixed tissue staining from the nPOD histopathology database show CD3+ infiltrated, insulin+ islets (**Figure 4g**).

### Beta Cell Glucose Metabolism is Disrupted During T1D Pathogenesis

Having established that remaining beta cells in slices from donors with T1D do not respond appropriately to glucose, and that this defect is not directly correlated with local T cell infiltrate, we set out to identify factors in beta cell glucose processing that could explain the dysfunction. Beta cells generate ATP upon supply of macronutrients resulting in insulin secretion. In brief, glucose enters human beta cells through glucose transporter 1 (GLUT1)^63^. Glucose is then phosphorylated by glucokinase followed by further metabolizing through glycolysis to generate pyruvate^64^. Pyruvate produced by glycolysis is transported by the mitochondrial pyruvate carriers where it is further metabolized to acetyl-CoA by pyruvate dehydrogenase. Acetyl-CoA enters the mitochondrial citric acid cycle leading to the production of reducing equivalents (NADH or FADH) that donate hydride for mitochondrial respiration and ATP synthesis. ATP synthesis and the subsequent decline in ADP levels increase the cytoplasmic ATP to ADP ratio. ATP displaces ADP in the ATP-sensitive K+ channels, leading to channel blockade and plasma membrane depolarization, and resultant increases in both action potential and intracellular Ca^2+ 64^. The increase in intracellular Ca^2+^ causes the readily releasable pool of insulin granules to fuse with the plasma membrane resulting in secretion.

To investigate this dysfunction further, we investigated changes in both RNA and protein expression involved in different facets of beta cell glucose metabolism from fixed pancreatic samples. We were able to access the large inventory of preserved nPOD specimens, we included additional donors in this analysis and were, thus, able to stratify sAAb+ and mAAb+ donor groups (see **Supplementary Table 1** for donor information). FFPE sections from ND (n=9), sAAb+ (n=7), mAAb+ (n=7), and T1D+ (n=6) donors were labeled with a panel of 22 antibodies (**Supplementary Table 3**) to investigate mechanisms underlying beta cell dysfunction in the main glucose metabolism pathway, including glucose import, glycolysis, oxidative phosphorylation, and membrane depolarization. Islet immunofluorescent (IF) confocal images of 30 insulin+ islets per donor were analyzed using Cell Profiler to allow measurement of marker MFI in only insulin+ cells^41^. Representative images from all markers in the panel are shown in **Supplementary Figure 7**. We also examined a separate cohort of laser-capture micro-dissected islets taken from frozen OCT sections from ND (n=10), sAAb+ (n=3), mAAb+ (n=3), and T1D+ donors (n=6) to evaluate changes in RNA expression for the same markers (**Supplementary Figure 7**).

Markers with significant differences in mean immunofluorescence intensity between insulin+ islets in ND and T1D+ donors – GLUT1, PFKFB4, GAPDH, SDH, HLA-ABC, KCNJ11, and CACNA1D – are shown alongside the insulin co-staining in the same islet (**Figure 5a**) and quantified in **Figure 5b**. As insulin secretion is the final step and critical outcome of the glucose metabolic pathway, we first investigated whether there were differences in insulin content as T1D progressed. RNA expression of whole laser-capture micro-dissected islets showed a decrease in *INS* RNA expression in T1D in the whole islet (**Figure 5c**), which may be expected with a decrease in beta cell content. However, when we investigated beta cells only within the insulin+ area of the IF-stained cohort, we found that insulin protein expression, on a per beta cell basis, remains constant across donor groups throughout the different stages of T1D development (**Figure 5b**, **Supplementary Figure 8**).

**Figure 5:**
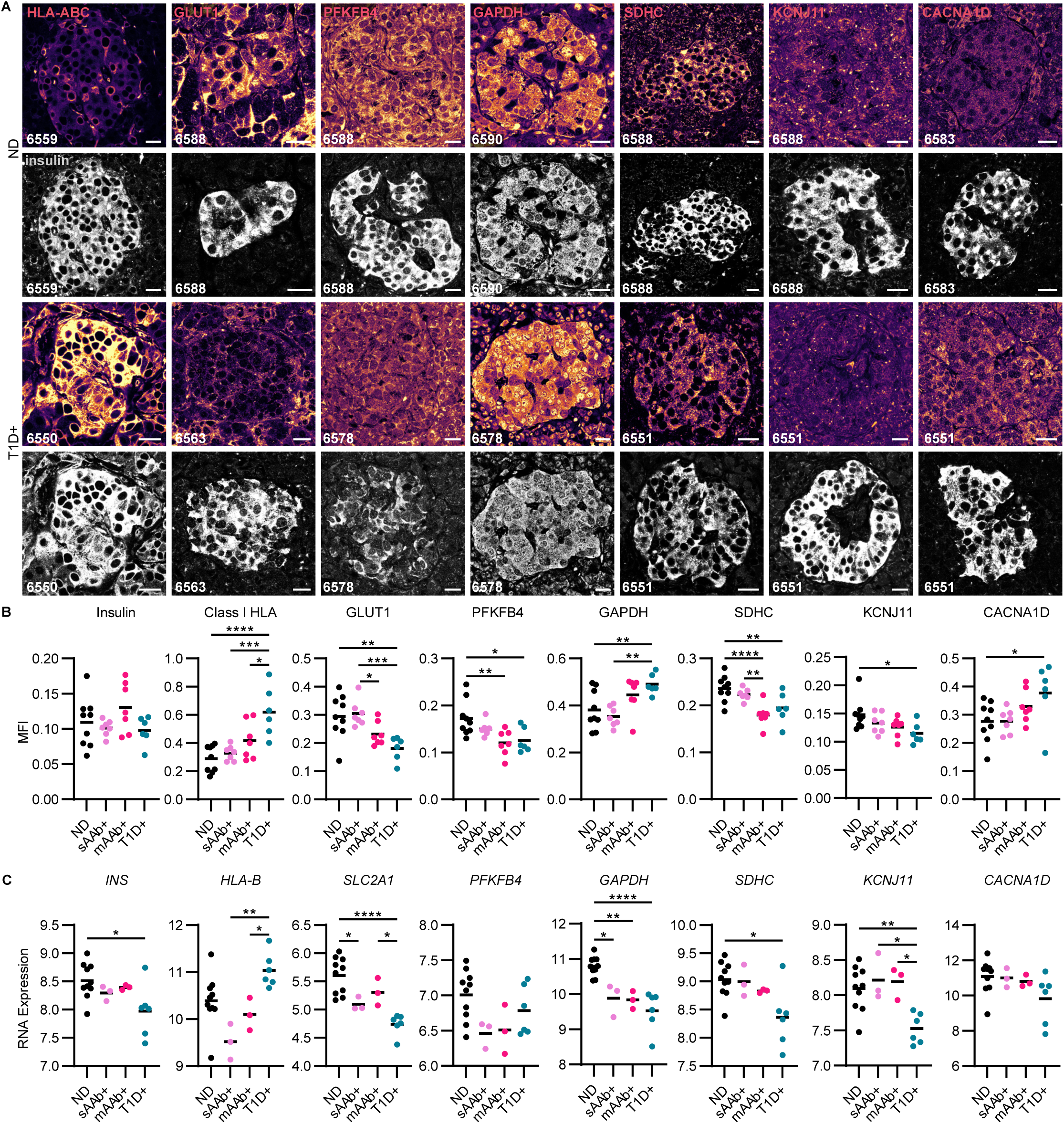
Beta cell glucose metabolism is disrupted during T1D pathogenesis. (**A**) Immunofluorescent images from ND and T1D+ donors representative of the dataset plotted in panel B. Scale bars, 20 µm. (**B**) Protein expression levels indicated by mean fluorescence intensity during the different stages of T1D development in donors with ND (n=9), sAab+ donors (n=7), mAab+ donors (n=7), and T1D+ donors (n=6). Scale bars, 20 µm. One-way ANOVA with multiple comparisons, *P<0.05, **P<0.01, ***P<0.001, ****P<0.0001. (**C**) RNA expression levels of glucose metabolisms markers in laser-capture micro-dissected islets from donors at different stages of T1D development (ND, n=10; sAab+, n=3; mAab+, n=3; T1D+, n=6). One-way ANOVA followed by a Tukey post-hoc test, *P<0.05, **P<0.01, ***P<0.001, ****P<0.0001.

The insulin staining validates that we included only insulin+ islet area in the MFI analysis and were not quantifying insulin-negative islets. We observed a strong upregulation of Class I HLA at both the protein level in the insulin+ islet area and the mRNA level in laser-captured whole islets from T1D+ donors (**Figure 5a-c**). This result is highly consistent with previous reports that the conditions of T1D pathogenesis result in increased islet visibility to the immune system through upregulation of class I HLA^65,66^.

The remaining markers with altered expression are directly involved in glucose metabolism. GLUT1 is the transporter responsible for glucose uptake into human beta cells^63^. GLUT1 staining was decreased in T1D+ donors (**Figure 5a,b**). *SLC2A1*, which encodes GLUT1, had decreased RNA expression in whole islets from sAAb+, mAAb+, and T1D+ donors (**Figure 5c**). We next studied PFKFB4, a regulator of the enzyme fructose-2,6-bisphosphate that acts as a rate-limiting step of glycolysis and as such, represents a critical step in glucose metabolism within the beta cell. Similar to GLUT1, PFKFB4 protein expression was also significantly decreased in mAAb+ and T1D+ donors (**Figure 5a,b**). The decrease in these critical glycolytic enzymes, not only in T1D+ but also mAAb+ donors, implies that a beta cell metabolic defect precedes widespread immune cell infiltration of islets.

GAPDH is the enzyme that catalyzes the sixth step of glycolysis converting glyceraldehyde 3-phosphate to 1,3-diphosphoglycerate. Within insulin+ cells, GAPDH had increased protein expression in T1D (**Figure 5a,b**) while there was downregulation of *GAPDH* RNA expression in whole laser-captured islets from sAAb+, mAAb+, and T1D+ donors (**Figure 5c**). To investigate this discrepancy, we visualized the subcellular localization of GAPDH within the beta cells as T1D progressed and found the increased GAPDH MFI in T1D was associated with bright nuclear staining that was absent in ND (**Figure 5a**). To contextualize this finding, it is critical to consider that in addition to serving as a glycolytic enzyme, GAPDH also plays a role in a cell death cascade, and as such, nuclear translocation of the protein indicates cellular stress^67–69^ and reduced catalytic activity^70^. We speculate that increased localization of GAPDH within the nucleus and overall downregulation of *GAPDH* mRNA are both consistent with poor glycolytic function.

Following glycolysis, the citric acid cycle and oxidative phosphorylation are the next major processes in beta cell glucose metabolism. As succinate dehydrogenase is involved in both processes, we next studied its subunit SDHC. MFI analysis of SDHC showed decreased expression in beta cells of both mAAb+ and T1D+ donors (**Figure 5a,b**). RNA expression of SDHC also decreased in T1D+ donors (**Figure 5c**). Again, this decrease present in mAAb+ as well as T1D+ donors implies a beta cell metabolic defect that occurs prior to islet-specific T cell infiltration. Several additional markers of mitochondrial function showed a reduction between ND and T1D at the mRNA level, but not the protein level, including *PC*, *OGDH*, *NDUFS1*, and *ATP5B* (**Supplementary Figure 6**).

Among proteins involved in membrane depolarization and insulin secretion, *KCNJ11* encodes the K_ir_6.2 subunit of the ATP-sensitive potassium (K_ATP_) channel. In beta cells, the K_ATP_ channel is essential for coupling glucose stimulation with insulin secretion. An increase in the [ATP]/[ADP] ratio from glucose metabolism closes the K_ATP_ channel, which depolarizes the membrane and consequently opens voltage-gated Ca^2+^ channels. KCNJ11 was significantly reduced at both the protein and mRNA levels in T1D+ donors, indicating a reduced capacity for membrane depolarization in insulin+ islets occurs in T1D prior to T cell infiltration (**Figure 5b,c**). Interestingly, the L-type calcium channel subunit Ca_v_1.3 encoded by *CACNA1D* was significantly upregulated at the protein but not mRNA level in T1D (**Figure 5b,c**). Increased voltage-gated Ca^2+^ channel expression may represent an attempt by the beta cell to compensate for decreased metabolic function or reduced channel turnover from overall lower activity.

As endoplasmic reticulum (ER) stress is frequently discussed as a possible contributor to beta cell death and dysfunction, we next investigated whether there were changes in expression of the stress markers IRE1α and PERK (**Supplementary Figure 6**) ^71–73^. IRE1α and PERK act as unfolded protein response stress sensors^74,75^. *ERN1*, which encodes IRE1α, had decreased RNA expression in sAAb+ and mAAb+ donors but not T1D. There were no differences in IRE1α protein expression between any of the donor groups. Whole islet RNA expression of *EIF2AK3*, which encodes PERK, revealed decreased expression in sAAb+, mAAb+, and T1D+ donors. However, similarly to protein expression of IRE1α, beta cell-specific PERK protein expression did not change between donor groups (**Supplementary Figure 6**). Overall, we found that the two ER stress markers we studied, IRE1α and PERK, did not change in expression as T1D progressed and thus, were not correlated with beta cell dysfunction.

We studied a total of 21 markers in the beta cell stimulus secretion coupling metabolic pathway by protein and RNA expression (**Figure 6a**). A heatmap was generated to visualize the protein expression intensity data using the ND group as the reference population and grouping markers by function. This overview of MFI changes highlights the reductions in beta cell proteins involved in glycolysis and mitochondrial respiration as T1D progresses (**Figure 6b**). Additionally, we assessed how the protein expression correlated with beta cell function and found protein expression of class I HLA, CACNA1D, GAPDH, SDHC, GCK, and GLUT1 all significantly correlated with the fraction of beta cells responsive to high glucose stimulation (**Supplementary Figure 8**).

**Figure 6:**
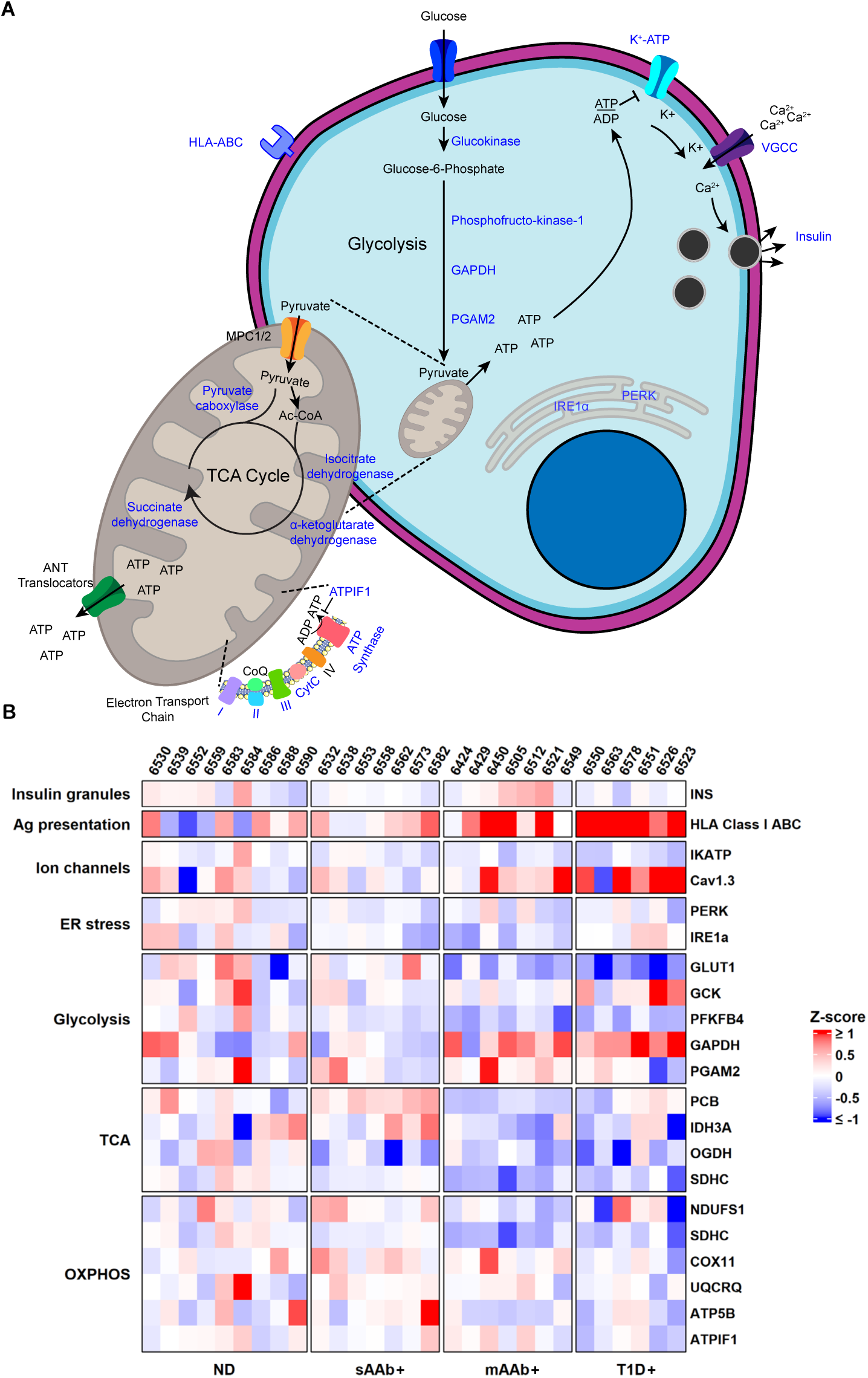
Downregulation of beta cell glucose metabolism markers occurs during T1D pathogenesis. (**A**) Schematic of beta cell glucose metabolism with the proteins targeted in these studies indicated in blue. (**B**) Heatmap visualization of protein expression. Red represents higher expression and blue represents lower expression relative to the ND group used as the baseline population for mean and standard deviation calculations. Proteins involved in glycolysis and mitochondrial respiration have expression changes as T1D pathogenesis progresses. The Ca^2+^ channel Cav1.3 and HLA Class I ABC also have increased expression in T1D+ donors.

## Discussion

The studies discussed herein have further established the period of beta cell dysfunction that occurs with T1D^7,8,10–14^. While many previous studies assessing beta cell function were conducted *in vivo* in patients, individuals at-risk for developing T1D, and healthy controls, the live pancreas slice model allows for this period of beta cell dysfunction to be investigated within the organ at a cellular level^7–16,27–30,39^.

Ca^2+^ recordings of slices from donors at different stages of disease development allow for investigations into what is occurring within the pancreas during T1D pathogenesis. Ca^2+^ recordings assess beta cell function at a smaller scale than perifusion experiments and provide a nuanced view of beta cell dysfunction. We applied the slice model to establish a loss of high glucose responses despite persistent maintenance of KCl responses in recordings of islets from T1D+ donors. These results clearly demonstrate that the beta cells within T1D+ tissues are not simply dead: they maintain a residual secretory capacity as demonstrated by the KCl responses yet no longer respond appropriately to glucose stimulation.

Studies using isolated human islets from patients with T1D have been extremely limited, but the published work does support the validity of our findings^47–50^. Immediately after isolation, islets from patients with T1D were deficient in GSIS. This loss of GSIS was not associated with a dearth of insulin expression but instead, with a significant reduction in the level of mRNA for members of glycolysis and the citric acid cycle when compared to islets isolated from individuals without T1D^47^. Using methods similar to those discussed above, for IF and gene expression, we also observed an increase in the inflammatory profile of T1D+ and mAAb+ islets compared to ND controls^76^. The previous studies, as well as our data, link the reductions in enzymes essential for GSIS with islet inflammation. Importantly, in the previous publications, long-term culture resulted in reductions of the inflammatory markers as well as the recovery of GSIS in islets isolated from organ donors with T1D^47,48^. When taken together, these results support our hypothesis that after the appearance of AAb, inflammatory signals act to downregulate the expression of genes essential for beta cell insulin secretory responses. The effects of these signals are reported to be reversible; therefore, elucidating the immune signals that negatively impact the beta cell will likely have a major impact on future treatment strategies^47–49^. Our study provides essential information on the function of beta cells from cases with T1D and at-risk organ donors in the presence or absence of autoreactive T cells.

The slice model is ideal for investigating beta cell dysfunction in T1D further because it allows for the simultaneous observation of beta cell functionality and T cell activity. This provided the unique opportunity to capture the first known recordings of human insulitis *in situ.* We also were able to detect how T cell infiltration correlated with beta cell function in T1D and found that T cells were not associated with worsening beta cell function. Conversely, beta cells in T1D+ slices exhibited dysfunctional responses to high glucose regardless of whether they were infiltrated by T cells. Thus, it appears that beta cell dysfunction is a defining feature of these cells in human T1D.

The next question we addressed was what beta cell defect(s) led to the loss of glucose metabolism observed during T1D development. We designed IF panels to assess protein expression changes during the critical steps of the glucose metabolism pathway in donors at different cross-sectional stages of T1D development. The most significant decreases at the protein level were found in glycolysis and the citric acid cycle. Defects were found to be present in mAAb+ and T1D+ donors. As both PFKFB4 and SDHC followed this trend, it is likely that beta cells are systemically downregulating components of glucose metabolism resulting in dysfunction. This, coupled with the increase of GAPDH nuclearization within the islet in T1D, emphasizes cellular stress and dysfunction. In summation, we have found that there are metabolic defects that precede disease diagnosis and that beta cells likely play a role in their own demise.

Our manuscript challenges the view that beta cell dysfunction in T1D must be closely associated, both spatially and temporally, with the local presence of insulitis. Rather, we show that beta cell dysfunction is not isolated to CD3+ islets and is consistent with other studies showing upregulation of Class I HLA ^66^, PD-L1 ^77^, and CXCL10 ^78^, widespread dysfunction of alpha cells, and exocrine atrophy throughout the pancreas in T1D ^79^. Prior studies that assessed beta cell function in human T1D came to differing conclusions on beta cell dysfunction in T1D. Studies performed in islets isolated from pancreata of donors with T1D generally concluded that glucose sensing and GSIS were preserved in the remaining beta cells when normalized to residual insulin content ^50,80,81^. Our interpretation is that these studies still demonstrated deficits in the first-phase insulin secretion and stimulation index relative to normal islets, which are hallmarks of T1D. Other studies showed that beta cells from donors with T1D are metabolically impaired when first isolated, but islets from a subset of donors recover GSIS when removed from the pancreatic microenvironment and cultured for several days under non-diabetic conditions ^48,49^.

Studies performed in human pancreas slices from donors with T1D differ dramatically from the studies in isolated islets. In slices, a profound and widespread loss of beta cell glucose sensing is observed that persists even when normalized to beta cell mass ^28,39^. Consistent with our hypothesis that beta cell impairment in T1D extends beyond the islets under direct immune attack, Panzer et al. found severe beta cell dysfunction even in regions of the pancreas that were devoid of T-cell infiltration ^28^.

The data in pancreas slices showing beta cell dysfunction in T1D contrasts with the studies in isolated islets. The discrepancies between these studies may be explained by different methodology (cultured islets vs. fresh pancreas slices), donor demographics (age and duration of diabetes), normalization approaches (insulin content vs. insulin-positive area), and interpretation of results (importance of first phase). Interestingly, these data suggest there are factors contributing to beta cell dysfunction present in the islet microenvironment of pancreas slices but not in isolated islets. Unlike isolated islets which is selective for those islets most likely to survive the isolation process, slices preserve the native pancreatic microenvironment, including small islets and islets with compromised structure, innate and adaptive immune cells, innervation, vasculature, and extracellular matrix all of which influence beta cell function and are influential in T1D etiopathology ^79^.

Throughout these studies, we have established the live pancreas tissue slice as a model to simultaneously assess beta cell function and immune cell infiltration. By combining this model and confocal microscopy, we were able to establish that beta cells are dysfunctional after T1D diagnosis regardless of T cell infiltration, a critical finding in T1D. We investigated the mechanisms of this dysfunction further by exploring changes in both the RNA and protein expression of the critical stages of glucose metabolism. We found decreases in glycolytic proteins as well as proteins involved in oxidative phosphorylation implying a systemic metabolic defect within the beta cell that precedes T1D diagnosis. These findings, coupled with the result that T cell infiltration does not correlate with beta cell dysfunction, indicate that systemic inflammation, as opposed to acute infiltration, likely initiates dysfunction. This is critical for understanding T1D pathogenesis and developing strategies to halt or slow its progression. Limitations of this study include heterogeneity of the donor population as is the case in studies in humans. Similarly, differences in donors’ health, age, cause of death, and cold ischemia time of the tissues should all be considered as well. Future directions include additional mechanistic studies to investigate these defects further in live functional tissues as opposed to fixed tissues to further establish the timeline and causes of this dysfunction and any possible resolutions.

## Acknowledgements

The authors thank the nPOD donors and their families for the gift of tissues, Stephan Speier (Technische Universität Dresden) and Denise Drotar (University of Florida) for assistance in initiating our work in human pancreas slices, and Andrece Powell (University of Florida) for technical assistance with immunostaining.

This work was funded by the following NIH grants, P01 AI042288 (EP, CM, MAA), R01 DK132387 (EP), R01 DK124267 (EP), R01 DK123292 (MA), R01 DK122160 (CM, MCT), U01 DK135001-01 (MCT), T32 DK108736 (MH), F31 DK130607 (MH), the NIDDK-supported Human Islet Research Network (HIRN, RRID:SCR_014393; https://hirnetwork.org; UH3 DK122638 (EP, CM), UC4 DK104155 (IG, CM, MCT), and R01 DK123329 (MCT)). Grant funds were also provided by Breakthrough T1D 2-SRA-2023-1313-S-B (EP). This research was also performed with the support of the Network for Pancreatic Organ donors with Diabetes (nPOD; RRID:SCR_014641), a collaborative type 1 diabetes research project supported by Breakthrough T1D and The Leona M. & Harry B. Helmsley Charitable Trust (Grant# 3-SRA-2023-1417-S-B). The content and views expressed are the responsibility of the authors and do not necessarily reflect the official view of nPOD. Organ Procurement Organizations (OPO) partnering with nPOD to provide research resources are listed at https://npod.org/for-partners/npod-partners/. This work utilized a Leica 7000 laser microdissection microscope purchased with an NIH shared instrumentation grant 1S10OD016350-01 and operated by the University of Florida Molecular Pathology Core (RRID:SCR_016601).

The beta cell schematic in Figure 5 was made using some graphics from Bioicons. Specifically, receptor-membrane-4 icon by Servier https://smart.servier.com/ is licensed under CC-BY 3.0 Unported https://creativecommons.org/licenses/by/3.0/, channel-membrane-lightpurple icon by Servier https://smart.servier.com/ is licensed under CC-BY 3.0 Unported https://creativecommons.org/licenses/by/3.0/, mitochondria icon by jaiganesh https://github.com/jaiganeshjg is licensed under CC0 https://creativecommons.org/publicdomain/zero/1.0/, Endoplasmic_Reticulum icon by jaiganesh https://github.com/jaiganeshjg is licensed under CC0 https://creativecommons.org/publicdomain/zero/1.0/, and emptycell-quarteroval-membrane-blue icon by Servier https://smart.servier.com/ is licensed under CC-BY 3.0 Unported https://creativecommons.org/licenses/by/3.0/

## Data availability statement

The RNA expression dataset is deposited at the Gene Expression Omnibus (GEO; accession number: GSE284772).

The 260 islets expression matrix is available from https://doi.org/10.5281/zenodo.14537115

The R code and analysis pipeline of the RNA expression dataset is posted to https://github.com/AlexandraCuaycal/laser_capture_microdissected_islet_transcriptomic s.

The R code and analysis pipeline of the Ca^2+^ imaging dataset is posted to https://github.com/PhelpsLabUF/Slice_Calcium_Imaging_Visualization_Analysis

The R code and analysis pipeline of the perifusion data analysis is posted to https://github.com/PhelpsLabUF/Perifusion_data_analysis.

The slice Ca^2+^ videos are available from https://doi.org/10.5281/zenodo.14605547

## Author Contributions

M.K.H conducted live human slice studies, performed fixed tissue staining and imaging, conducted Ca^2+^ imaging analysis, fixed tissue analysis, and drafted the manuscript. A.E.W. performed the R analysis of the Ca^2+^ imaging and fixed tissue imaging data. A.E.C. performed the R analysis of the RNA expression dataset and the perifusion data. D.S. performed T cell tracking and analysis. E.A.B. and M.C.T. conducted islet microdissection, RNA isolation, and slide staining and analyses. J.C participated in islet microdissection. I.C.G. conducted microarray analysis. N.I.L. analyzed global gene expression patterns in RNA samples using microarrays and conducted higher level data analysis. M.B., H.H., generated live human pancreas tissue slices and conducted perifusion studies. E.V. generated live human pancreas tissue slices. I.K. coordinated nPOD organ procurement, tissue processing, slice generation, and fixed tissue slide generation. M.S.R. advised on Ca^2+^ imaging protocols and analysis. M.C.T. provided guidance on fixed tissue staining and use of equipment and lab space for staining. M.A.A. acquired funding, supervised research, and advised on studies. C.E.M and E.A.P. acquired funding, conceived of, supervised and provided guidance on the studies, and edited the manuscript. E.A.P. developed the approaches for imaging of insulitic islets from T1D donors. C.E.M. developed the hypothesis that was tested. All authors discussed the results, commented on the manuscript, and approved of its submission.

---

**Supplementary Figure 1:**
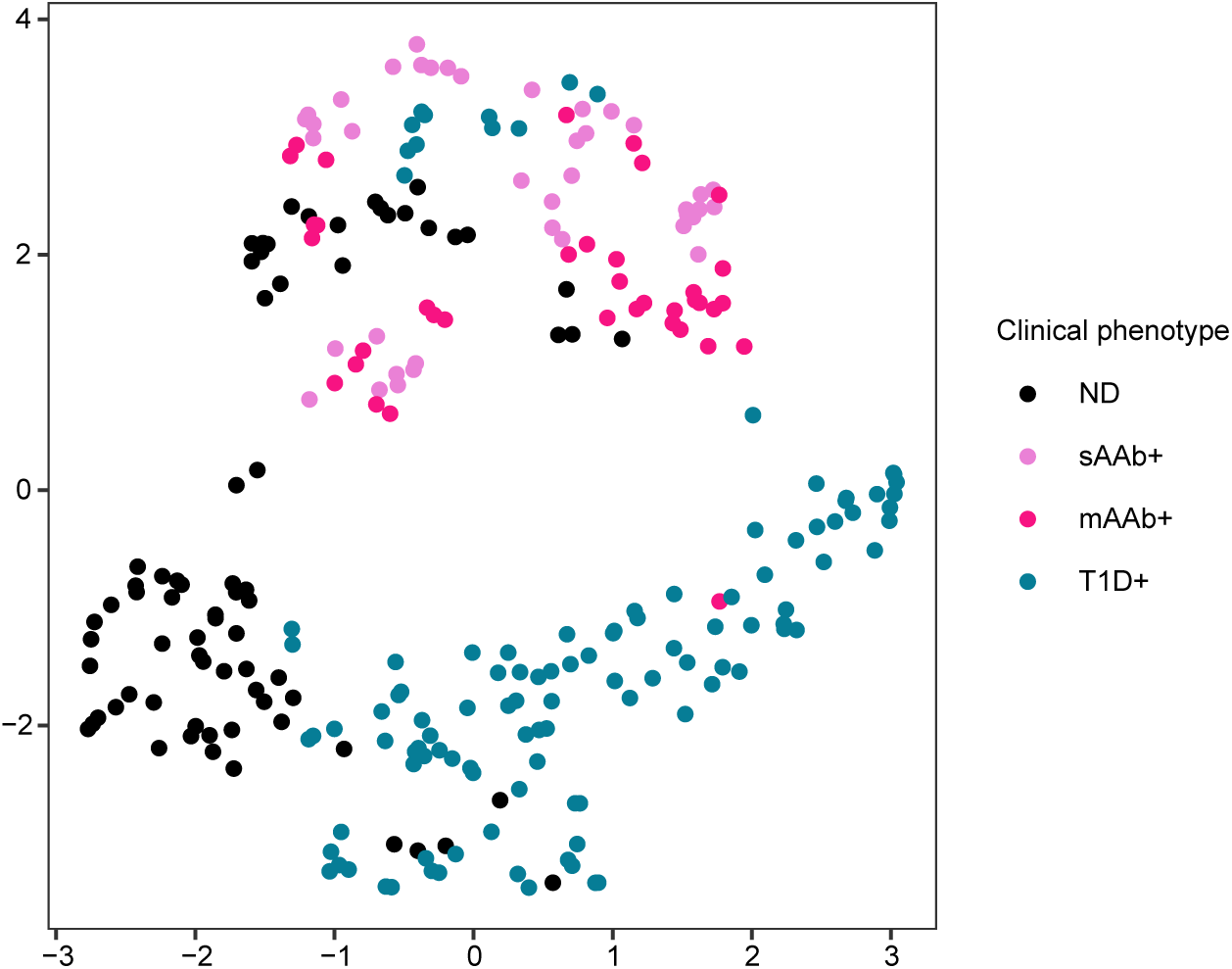
UMAP of laser-capture microdissected islets. Uniform manifold approximation and projection (UMAP) plot showing clustering of 260 islets. Each dot represents an islet and each clinical phenotype is plotted with different color.

**Supplementary Figure 2:**
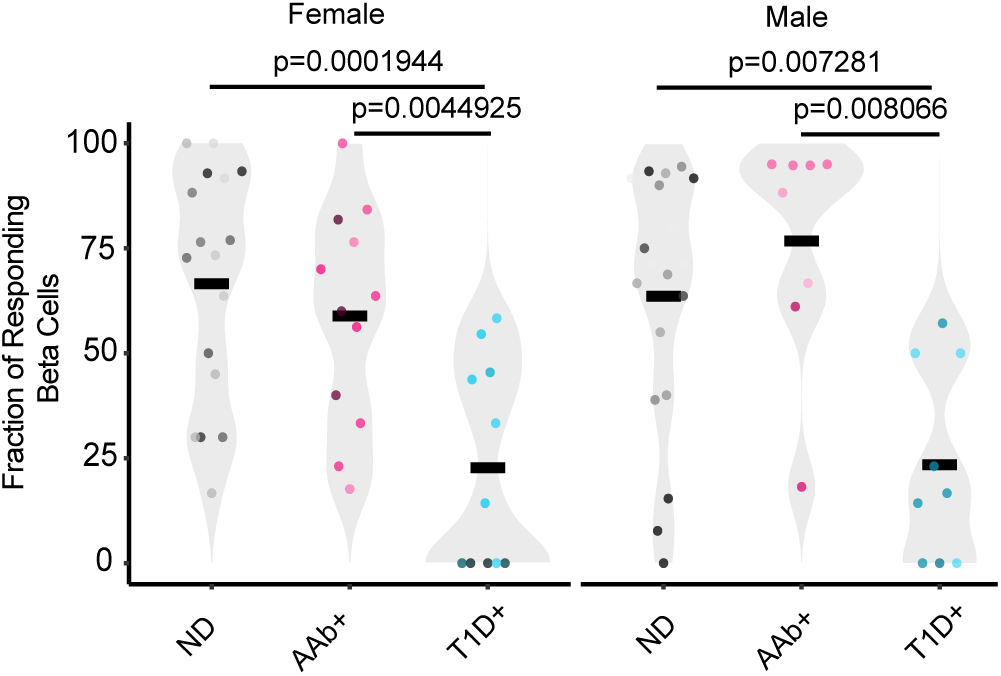
Beta cell dysfunction is not driven by sex differences. The fraction of beta cells responding to high glucose within each islet in male and female donors recorded from ND, Aab+, and T1D+ cases. Each dot represents an islet recording. Center line indicates the mean. One-way ANOVA with multiple comparisons.

**Supplementary Figure 3:**
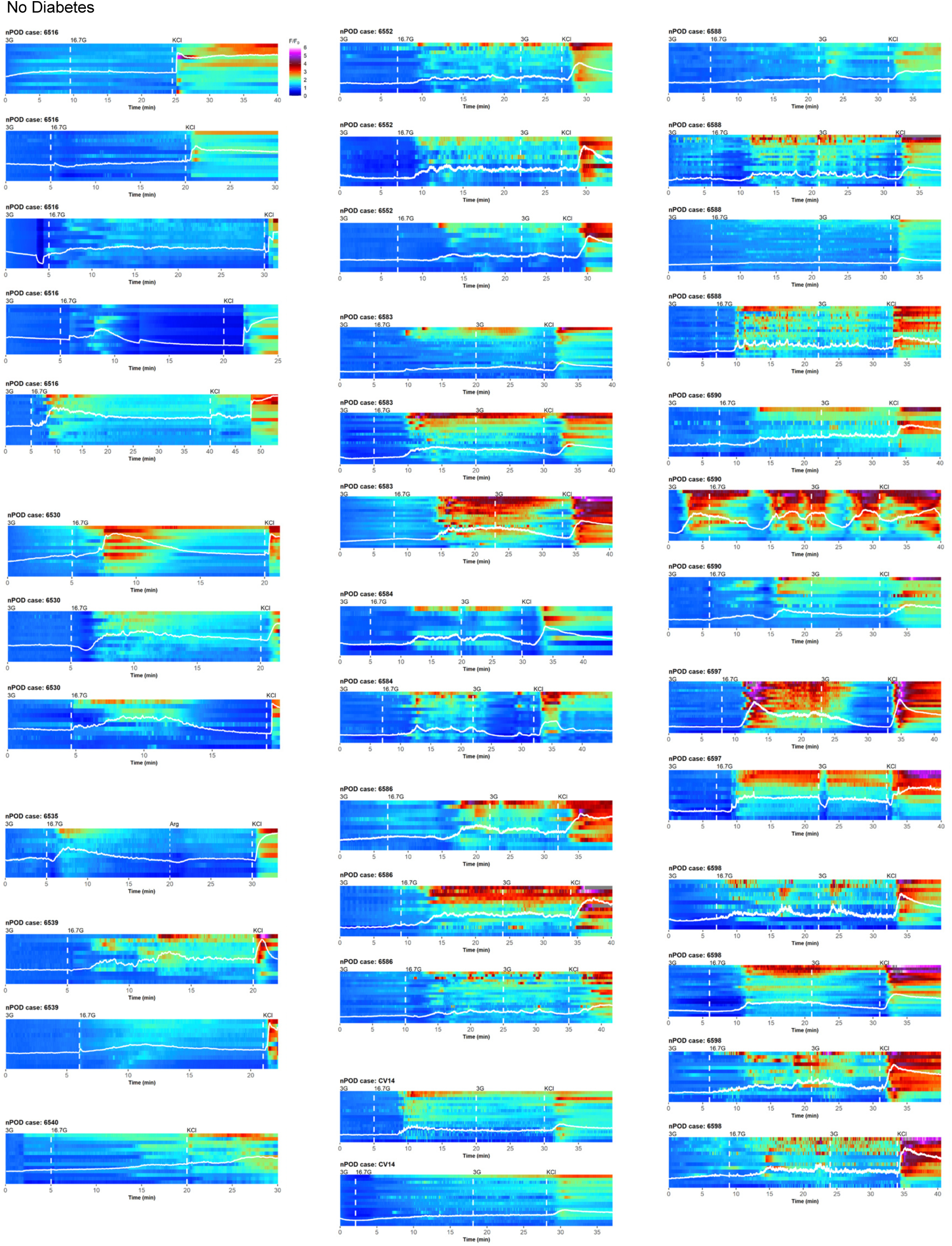
Ca^2+^ responses of beta cells within ND slices. Heatmaps of all ND Ca^2+^ recordings showing the changes in fluorescence of the individual ROIs from ND cases along with the superimposed average traces showing the glucose and KCl responses.

**Supplementary Figure 4:**
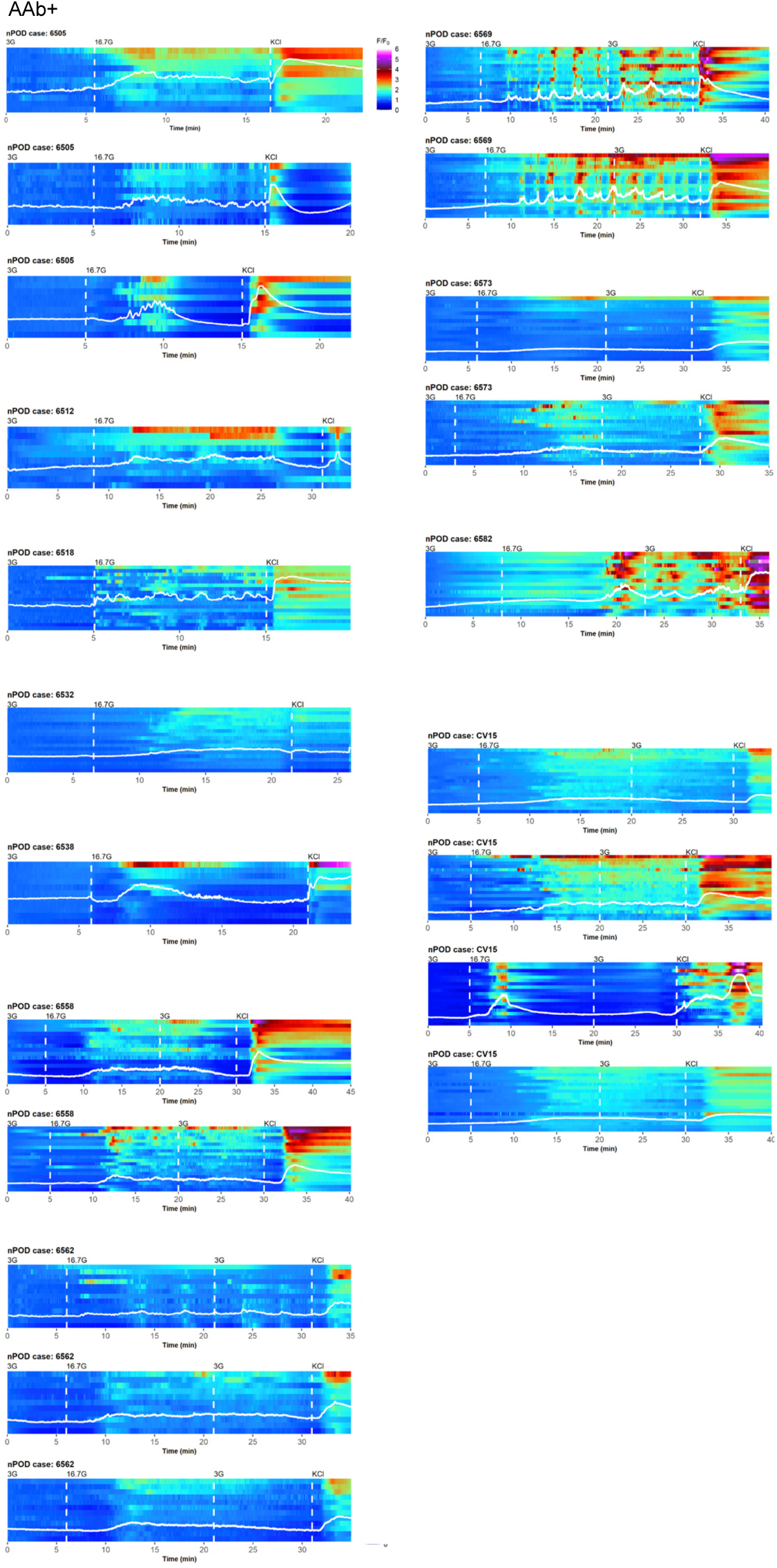
Ca^2+^ responses of beta cells within Aab+ slices. Heatmaps of all Aab+ Ca^2+^ recordings showing the changes in fluorescence of the individual ROIs from Aab+ cases along with the superimposed average traces showing the glucose and KCl responses.

**Supplementary Figure 5:**
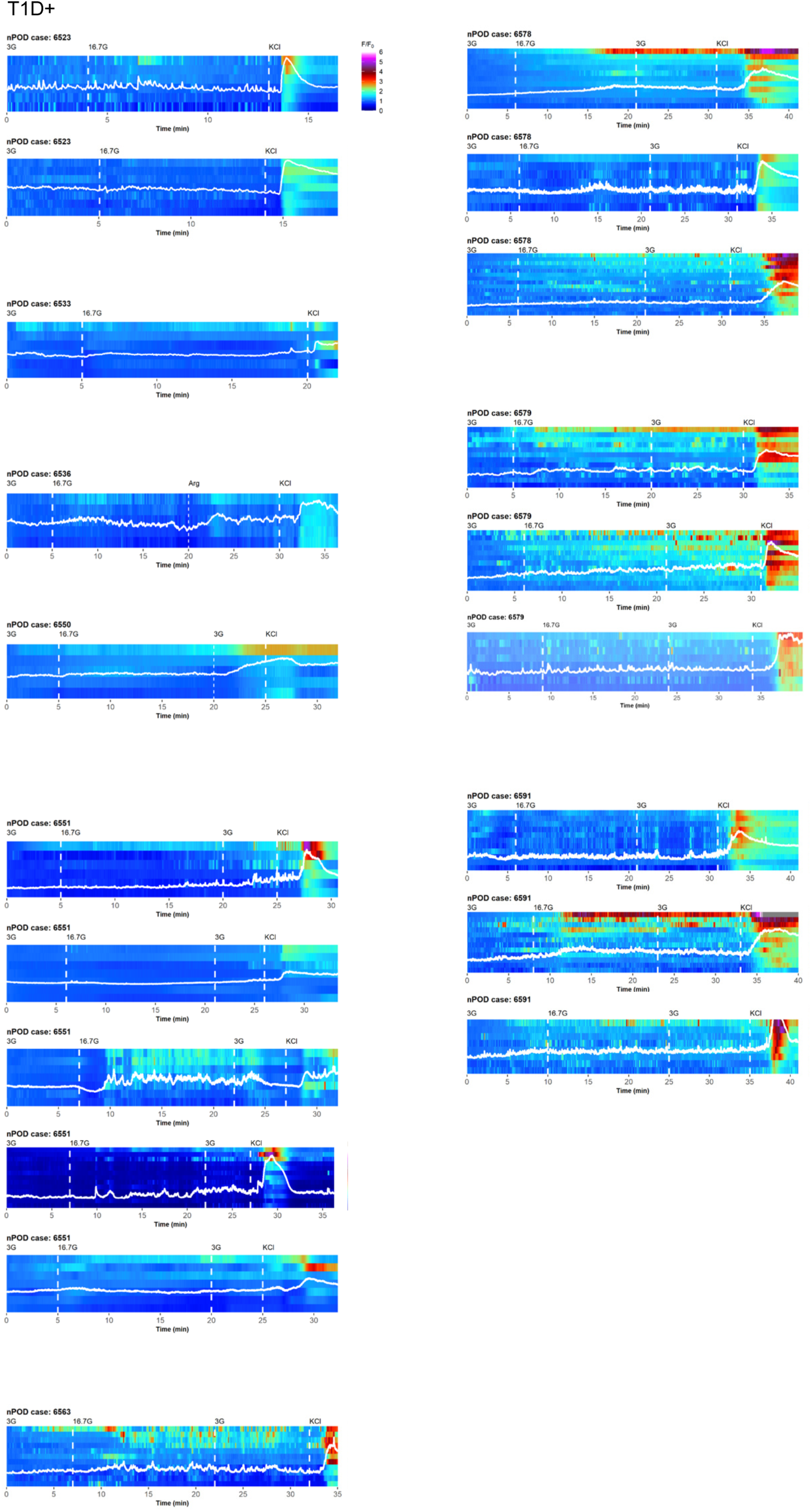
Ca^2+^ responses of beta cells within T1D+ slices. Heatmaps of all T1D+ Ca^2+^ recordings showing the changes in fluorescence of the individual ROIs from T1D+ cases along with the superimposed average traces showing the lost high glucose responses and maintained KCl responses.

**Supplementary Figure 6:**
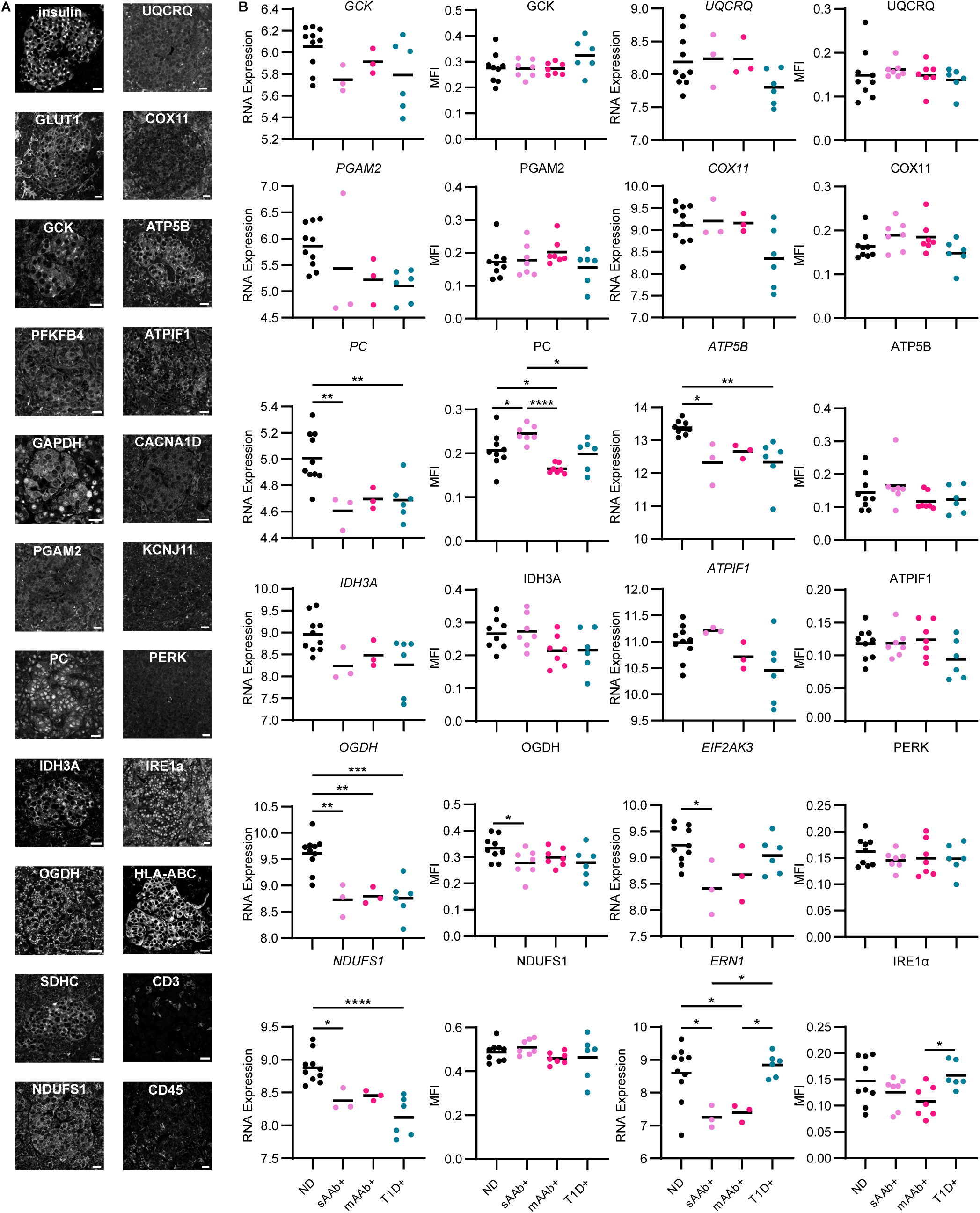
ER stress markers and some glucose metabolism markers do not differ during T1D pathogenesis. (**A**) Representative images of immunofluorescent staining of FFPE slides of the panel of markers used to assess immune cell infiltration and beta cell glucose metabolism in donors with ND (n=9), sAab+ donors (n=7), mAab+ donors (n=7), T1D+ donors (n=6). Scale bars, 20 µm. (**B**) RNA expression levels of glucose metabolisms markers in laser-capture micro-dissected islets from donors at different stages of T1D development (ND, n=10; sAab+, n=3; mAab+, n=3; T1D+, n=6) and protein expression levels indicated by mean fluorescence intensity during the different stages of T1D development in donors with ND (n=9), sAab+ donors (n=7), mAab+ donors (n=7), and T1D+ donors (n=6). MFIs: One-way ANOVA with multiple comparisons, *P<0.05, **P<0.01, ***P<0.001, ****P<0.0001. RNA: One-way ANOVA followed by a Tukey post-hoc test, *P<0.05, **P<0.01, ***P<0.001, ****P<0.0001.

**Supplementary Figure 7:**
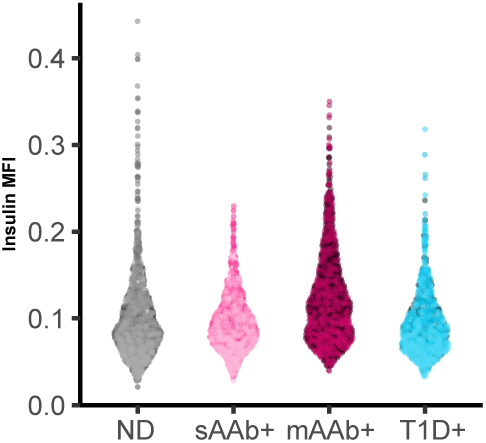
Insulin protein levels within beta cells does not change as T1D progresses. Protein expression levels of insulin within beta cells indicated by mean fluorescence intensity during the different stages of T1D development in donors with ND (n=9), sAab+ donors (n=7), mAab+ donors (n=7), and T1D+ donors (n=6). Each dot represents an islet. One-way ANOVA with multiple comparisons,

**Supplementary Figure 8:**
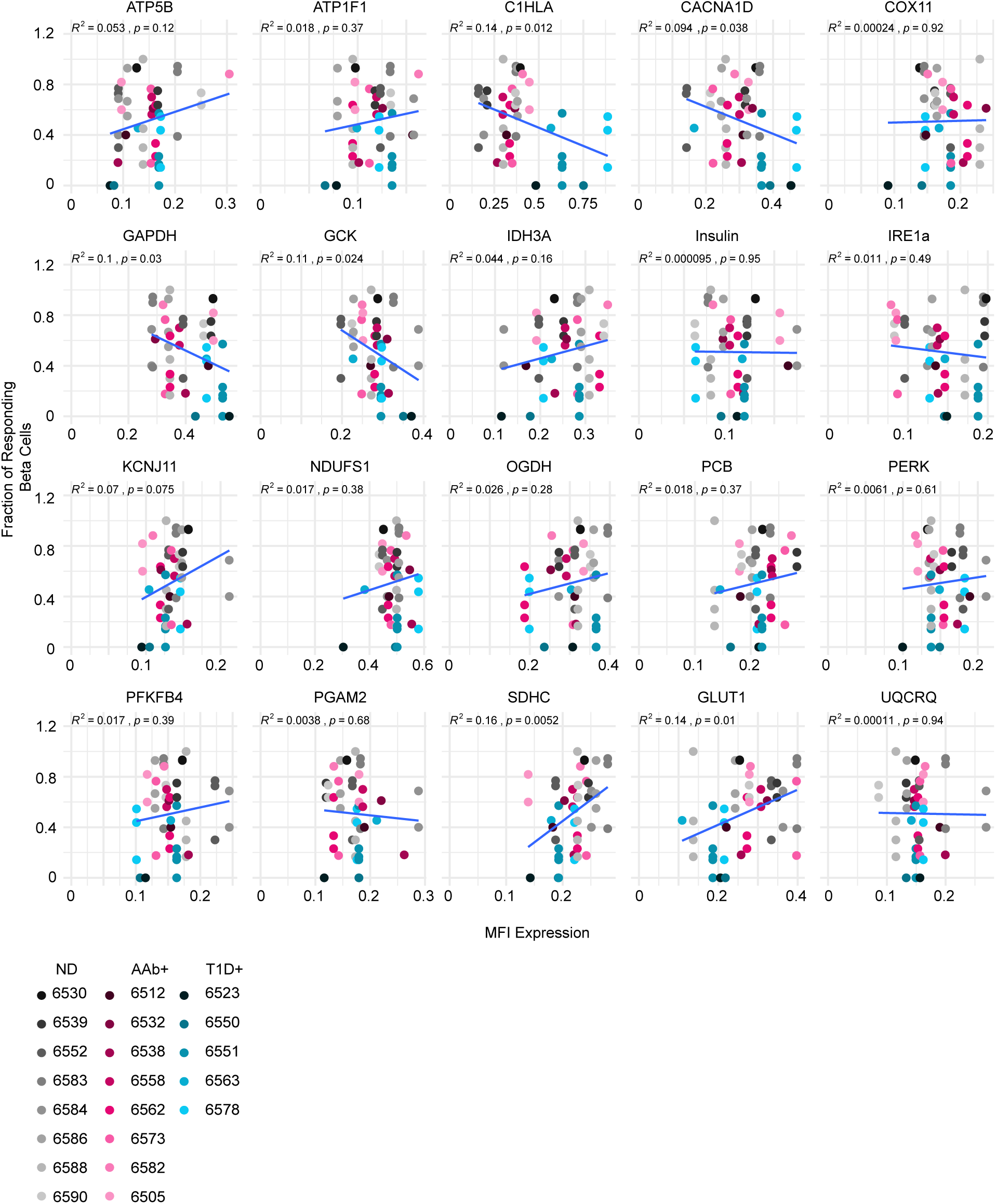
Correlation of beta cell function and glucose metabolism markers. Graphs showing the relationship between the fraction of responding beta cells and protein expression of markers of glucose metabolism with the linear models indicated in blue.

**Supplementary Table 1:** Donor characteristics from nPOD cases. Table summarizes donor information and applied studies.

| Case | RRID | Type | Duration (years) | Autoantibody | Age (years) | Sex | Ethnicity | HbA1c | C-pep (ng/mL) | HLA-DR | HLA-DQ | Data Types |
| --- | --- | --- | --- | --- | --- | --- | --- | --- | --- | --- | --- | --- |
| CV14 | SAMN38117313 | No Diabetes |  | Negative | 9.85 | F | Caucasian | 5.3 | 4.37 | DR07/15 | DQ02/06 | Ca<sup>2+</sup> recordings, perfusion |
| 6073 | SAMN15879130 | No Diabetes |  | Negative | 19.2 | M | Caucasian | 0 | 0.69 | DR13/16 | DQ05/06 | RNA expression |
| 6098 | SAMN15879155 | No Diabetes |  | Negative | 17.8 | M | Caucasian | 4.9 | 1.41 | DR17/08 | DQ02/04 | RNA expression |
| 6131 | SAMN15879188 | No Diabetes |  | Negative | 24.2 | M | Caucasian | 0 | 1.01 | DR17/17 | DQ02/02 | RNA expression |
| 6230 | SAMN15879286 | No Diabetes |  | Negative | 16 | M | Caucasian | 5.3 | 5.22 | DR04/11 | DQ07/08 | RNA expression |
| 6235 | SAMN15879291 | No Diabetes |  | Negative | 30 | M | Caucasian | 0 | 8.1 | DR04/17 | DQ02/08 | RNA expression |
| 6293 | SAMN15879347 | No Diabetes |  | Negative | 9 | F | Caucasian | 0 | 2.22 | DR01/11 | DQ05/07 | RNA expression |
| 6316 | SAMN15879370 | No Diabetes |  | Negative | 6 | M | Caucasian | 0 | 4.58 | DR04/13 | DQ08/09 | RNA expression |
| 6336 | SAMN15879390 | No Diabetes |  | Negative | 14.3 | F | Caucasian | 5.2 | 7.87 | DR07/13 | DQ02/06 | RNA expression |
| 6339 | SAMN15879393 | No Diabetes |  | Negative | 23.3 | M | Caucasian | 5.3 | 10.56 | DR03/10 | DQ02/05 | RNA expression |
| 6401 | SAMN15879454 | No Diabetes |  | Negative | 25.07 | F | Hispanic/ Latino | 5.8 | 12.81 | DR07/13 | DQ02/06 | RNA expression |
| 6516 | SAMN18053200 | No Diabetes |  | Negative | 20.75 | M | Caucasian | 5.5 | 8.91 | DR01/07 | DQ02/05 | Ca<sup>2+</sup> recordings, perfusion |
| 6530 | SAMN18053212 | No Diabetes |  | Negative | 14.52 | F | African American | 6 | 7.41 | DR04/17 | DQ02/08 | Ca<sup>2+</sup> recordings, IF, perfusion |
| 6535 | SAMN18242779 | No Diabetes |  | Negative | 31.07 | F | Caucasian |  | 6.59 | DR11/13 | DQ06/07 | Ca<sup>2+</sup> recordings |
| 6539 | SAMN25652250 | No Diabetes |  | Negative | 24.6 | M | Hispanic | 5.7 | 39.23 | DR04/13 | DQ06/08 | Ca<sup>2+</sup> recordings, IF |
| 6540 | SAMN25652251 | No Diabetes |  | Negative | 6.86 | F | Hispanic | 5.8 | 2.03 | DR04/11 | DQ07/04 | Ca<sup>2+</sup> recordings, perfusion |
| 6552 | SAMN30386842 | No Diabetes |  | Negative | 33.87 | F | Caucasian | 5.6 | 1.8 | DR17/04 | DQ02/08 | Ca<sup>2+</sup> recordings, IF, perfusion |
| 6559 | SAMN30386848 | No Diabetes |  | Negative | 23.46 | F | Asian | 5.3 | 7.02 | DR11/12 | DQ07/-- | IF |
| 6583 | SAMN38117303 | No Diabetes |  | Negative | 30 | M | Caucasian | 5 | 10.55 | DR11/13 | DQ07/06 | Ca<sup>2+</sup> recordings, IF, perfusion |
| 6584 | SAMN38117304 | No Diabetes |  | Negative | 22 | M | Caucasian | 5.3 | 6.02 | DR04/11 | DQ07/08 | Ca<sup>2+</sup> recordings, IF, perfusion |
| 6586 | SAMN38117306 | No Diabetes |  | Negative | 13 | M | Caucasian | 5.1 | 6.83 | DR04/07 | DQ02/08 | Ca<sup>2+</sup> recordings, IF, perfusion |
| 6588 | SAMN40555580 | No Diabetes |  | Negative | 29 | F | Caucasian | 5.1 | 7.3 | DR04/11 | DQ07/07 | Ca<sup>2+</sup> recordings, IF, perfusion |
| 6590 | SAMN38117309 | No Diabetes |  | Negative | 16 | F | African American | 5.3 | 49.65 | DR08/13 | DQ06/07 | Ca<sup>2+</sup> recordings, IF, perfusion |
| 6597 | SAMN40555584 | No Diabetes |  | Negative | 6 | F | Hispanic/Latino | 5.7 | 5.02 | DR04/04 | DQB08/08 | Ca<sup>2+</sup> recordings |
| 6598 | SAMN40555585 | No Diabetes |  | Negative | 17 | M | Hispanic/Latino | 5.4 | 1.36 | DR04/14 | DQB07/08 | Ca<sup>2+</sup> recordings |
| CV15 | SAMN40555609 | AAb+ |  | GADA+ | 18.93 | M | Caucasian | 5.2 | 5.03 | DR07/15 | DQ06/09 | Ca<sup>2+</sup> recordings, perfusion |
| 6080 | SAMN15879137 | AAb+ |  | GADA+, mIAA+ | 69.2 | F | Caucasian | 0 | 1.84 | DR01/04 | DQ03/05 | RNA expression |
| 6147 | SAMN15879203 | AAb+ |  | GADA+ | 23.8 | F | Caucasian | 5.2 | 3.19 | DR04/08 | DQ04/07 | RNA expression |
| 6158 | SAMN15879214 | AAb+ |  | GADA+, mIAA+ | 40.3 | M | Caucasian | 5.6 | 0.51 | DR04/13 | DQ06/07 | RNA expression |
| 6167 | SAMN15879223 | AAb+ |  | IA2A+, ZnT8A+ | 37 | M | Caucasian | 0 | 5.43 | DR04/02 | DQ08/06 | RNA expression |
| 6181 | SAMN15879237 | AAb+ |  | GADA+ | 31.9 | M | Caucasian | 0 | 0.06 | DR01/04 | DQ08/05 | RNA expression |
| 6314 | SAMN15879368 | AAb+ |  | GADA+ | 21 | M | Caucasian | 0 | 1.49 | DR103/04 | DQ03/05 | RNA expression |
| 6424 | SAMN15879477 | AAb+ |  | GADA+, mIAA+ | 17.65 | M | Caucasian | 5.8 | 6.97 | DR17/04 | DQ02/08 | IF |
| 6429 | SAMN15879482 | AAb+ |  | GADA+, mIAA+ | 22.1 | M | African American | 5.5 | 2.25 | DR103/17 | DQ02/05 | IF |
| 6450 | SAMN15879503 | AAb+ |  | GADA+, ZnT8A+ | 22 | F | Caucasian | 5.7 | 5.47 | DR17/-- | DQ02/-- | IF |
| 6505 | SAMN15879558 | AAb+ |  | GADA+, mIAA+ | 20.59 | F | Hispanic | 5.1 | 20.8 | DR04/07 | DQ02/08 | Ca<sup>2+</sup> recordings, IF, perfusion |
| 6512 | SAMN15879564 | AAb+ |  | IA2A+, mIAA+, ZnT8A+ | 30.59 | F | Caucasian | 5.2 | 3.2 | DR04/07 | DQ02/07 | Ca<sup>2+</sup> recordings, IF, perfusion |
| 6518 | SAMN25652247 | AAb+ |  | GADA+ | 21.86 | M | Caucasian | 6 | 10.93 | DR01/01:03 | DQ05/05 | Ca<sup>2+</sup> recordings, perfusion |
| 6521 | SAMN18053204 | AAb+ |  | GADA+, IA2A+, ZnT8A+ | 19.77 | M | Hispanic/ Latino | 5.8 | 7.44 | DR04/17 | DQ02/08 | IF |
| 6532 | SAMN18053214 | AAb+ |  | GADA+ | 20.04 | M | Hispanic | 5.9 | 22.12 | DR07/11 | DQ02/07 | Ca<sup>2+</sup> recordings, IF, perfusion |
| 6538 | SAMN25652249 | AAb+ |  | GADA+ | 19.14 | M | Caucasian | 5.8 | 11.33 | DR17/08 | DQ02/04 | Ca<sup>2+</sup> recordings, IF, perfusion |
| 6549 | SAMN25652260 | AAb+ |  | GADA+, mIAA+ | 4.22 | M | Caucasian | 6.2 | 1.17 | DR04/07 | DQ08/09 | IF |
| 6553 | SAMN30386843 | AAb+ |  | mIAA+ | 12.34 | F | Hispanic/ Latino | 8.4 | 4.62 | DR17/04 | DQ02/08 | IF |
| 6558 | SAMN30386847 | AAb+ |  | GADA+ | 21.69 | F | African American | 4.4 | 8.03 | DR17/07 | DQ02/-- | Ca<sup>2+</sup> recordings, IF |
| 6562 | SAMN30386850 | AAb+ |  | GADA+ | 29.77 | F | Caucasian | 5.6 | 13.19 | DR04/15 | DQ06/08 | Ca<sup>2+</sup> recordings, IF |
| 6569 | SAMN38117299 | AAb+ |  | GADA+ | 20 | F | Hispanic | 6 | 2.44 | DR04/15 | DQ08/06 | Ca<sup>2+</sup> recordings, perfusion |
| 6573 | SAMN33284291 | AAb+ |  | GADA+ | 24.48 | F | Caucasian | 5.1 | 1.85 | DR01/17 | DQ02/05 | Ca<sup>2+</sup> recordings, IF, perfusion |
| 6582 | SAMN38117302 | AAb+ |  | GADA+ | 22.38 | M | Caucasian | 5.6 | 2.67 | DR17/04 | DQ02/08 | Ca<sup>2+</sup> recordings, IF, perfusion |
| 6268 | SAMN15879322 | T1D+ | 3 | mIAA+ | 12 | F | Caucasian | 9.8 | 0.05 | DR13/17 | DQ02/06 | RNA expression |
| 6306 | SAMN15879360 | T1D+ | 5 | mIAA+ | 19 | M | Caucasian | 10.1 | <0.02 | DR04/17 | DQ02/08 | RNA expression |
| 6342 | SAMN15879396 | T1D+ | 2 | IA2A+, mIAA+ | 14 | F | Caucasian | 9.2 | 0.26 | DR01/04 | DQ08/05 | RNA expression |
| 6362 | SAMN15879415 | T1D+ | 0 | GADA+ | 24.9 | M | Caucasian | 10 | 0.38 | DR103/17 | DQ02/05 | RNA expression |
| 6371 | SAMN15879424 | T1D+ | 2 | GADA+, IA2A+, mIAA+, ZnT8A+ | 12.5 | F | Caucasian | 9.5 | 0.11 | DR13/17 | DQ02/06 | RNA expression |
| 6396 | SAMN15879449 | T1D+ | 2 | Negative | 17.1 | F | Caucasian | 13.4 | 0.06 | DR07/17 | DQ02/null | RNA expression |
| 6523 | SAMN18053206 | T1D+ | 3 | GADA+, mIAA+ | 12.17 | F | African American | 11.1 | 0.04 | DR17/07 | DQ02/02 | Ca<sup>2+</sup> recordings, IF |
| 6526 | SAMN18053209 | T1D+ | 1 | GADA+, mIAA+ | 29.74 | M | Hispanic/ Latino | 6.6 | 0.07 | DR04/13 | DQ08/06 | IF |
| 6533 | SAMN18242777 | T1D+ | 0 | IA2A+, mIAA+, ZnT8A+ | 3.75 | F | Caucasian | 11.4 | 0.17 | DR04/17 | DQ02/08 | Ca<sup>2+</sup> recordings, perfusion |
| 6536 | SAMN18242780 | T1D+ | 4 | GADA+ | 20.16 | F | Caucasian | 12.7 | 0.04 | DR04/17 | DQ02/08 | Ca<sup>2+</sup> recordings, perfusion |
| 6550 | SAMN25652261 | T1D+ | 0 | GADA+, ZnT8A+ | 25.06 | M | Caucasian | 14 | 0.02 | DR17/-- | DQ02/07 | Ca<sup>2+</sup> recordings, IF, perfusion |
| 6551 | SAMN25652262 | T1D+ | 0.58 | GADA+, IA2A+, mIAA+, ZnT8A+ | 20.7 | M | Caucasian | 6.4 | 0.11 | DR04/07 | DQ02/07 | Ca<sup>2+</sup> recordings, IF, perfusion |
| 6563 | SAMN30386851 | T1D+ | 0 | IA2A+ | 14.56 | F | Caucasian | 9.6 | 1.04 | DR07/08 | DQ02/04 | Ca<sup>2+</sup> recordings, IF, perfusion |
| 6578 | SAMN33284295 | T1D+ | 0 | IA2A+, ZnT8A+ | 11.95 | F | Caucasian | 13.6 | 0.35 | DR04/11 | DQ07/08 | Ca<sup>2+</sup> recordings, IF, perfusion |
| 6579 | SAMN33284296 | T1D+ | 1 | GADA+, mIAA+ | 13 | F | Caucasian | 15 | 0.31 | DR17/13 | DQB02/06 | Ca<sup>2+</sup> recordings, perfusion |
| 6591 | SAMN38117310 | T1D+ | 0 | IA2A+, ZnT8A+ | 3 | M | Caucasian | 12.8 | 0.03 | DR04/13 | DQ08/06 | Ca<sup>2+</sup> recordings, perfusion |

**Supplementary Table 2:** Ca^2+^ imaging summary. Ca^2+^ imaging characteristics and protocols for all recordings.

| Case | Cell Selection | Slice | Ca<sup>2+</sup> Indicator | Protocol | Frame Rate (s) |
| --- | --- | --- | --- | --- | --- |
| 6505 | Reflectance | 2 | Fluo4 | 0-5 min LG, 5-15 min HG, 15 min-end KCl | 2.873 |
| 6505 | Reflectance | 5 | Fluo4 | 0-5 min LG, 5-15 min HG, 15 min-end KCl | 1.99913 |
| 6505 | Reflectance | 6 | Fluo4 | 0-5 min LG, 5-15 min HG, 16 min-end KCl | 1.99886 |
| 6512 | Reflectance | 2 | Fluo4 | 0-7 min LG, 7-22 min 30s HG, 22 min 30s -end KCl | 1.99986 |
| 6516 | Reflectance | 1 | Fluo4 | 0-5 min LG, 5-20 min HG, 20 min-end KCl | 2.31722 |
| 6516 | Reflectance | 4 | Fluo4 | 0-5 min LG, 5-20 min HG, 20 min-end KCl | 2.31152 |
| 6516 | Reflectance | 6 | Fluo4 | 0-5 min LG, 5-30 min HG, 30 min-end KCl | 2.31424 |
| 6516 | Reflectance | 7 | Fluo4 | 0-5 min LG, 5-20 min HG, 20 min-end KCl | 2.31447 |
| 6516 | Reflectance | 8 | Fluo4 | 0-5 min LG, 5-40 min HG, 40 min-end KCl | 2.31152 |
| 6518 | Reflectance | 4 | Fluo4 | 0-5 min LG, 5-15 min HG, 15 min-end KCl | 3.52091 |
| 6523 | Reflectance | 3 | Fluo4 | 0-3 min LG, 3-18 min HG, 18 min-end KCl | 2.72597 |
| 6523 | Reflectance | 4 | Fluo4 | 0-3 min LG, 3-18 min HG, 18 min-end KCl | 2.72598 |
| 6530 | ENTPD3 | 1 | Fluo4 | 0-5 min LG, 5-20 min HG, 20 min-end KCl | 1.99799 |
| 6530 | ENTPD3 | 3 | Fluo4 | 0-5 min LG, 5-20 min HG, 20 min-end KCl | 1.99883 |
| 6530 | ENTPD3 | 4 | Fluo4 | 0-5 min LG, 5-20 min HG, 20 min-end KCl | 1.99834 |
| 6532 | ENTPD3 | 6 | Fluo4 | 0-5 min LG, 5-20 min HG, 20 min-end KCl | 1.99933 |
| 6533 | ENTPD3 | 10 | Fluo4 | 0-5 min LG, 5-20 min HG, 20 min-end KCl | 2.55227 |
| 6535 | ENTPD3 | 8 | Fluo4 | 0-5 min LG, 5-20 min HG, 20-30 min arginine, 30 min-end KCl | 2.31722 |
| 6536 | ENTPD3 | 4 | Fluo4 | 0-5 min LG, 5-20 min HG, 20-30 min arginine, 30 min-end KCl | 2.31721 |
| 6538 | ENTPD3 | 4 | Fluo4 | 0-5 min 50s LG, 5 min 50s- 21 min HG, 21 min-end KCl | 2.31696 |
| 6539 | ENTPD3 | 1 | Fluo4 | 0-5 min LG, 5-20 min HG, 20 min-end KCl | 2.31335 |
| 6539 | ENTPD3 | 2 | Fluo4 | 0-5 min LG, 5-20 min HG, 20 min-end KCl | 2.31407 |
| 6540 | ENTPD3 | 3 | Fluo4 | 0-5 min LG, 5-20 min HG, 20 min-end KCl | 2.31721 |
| 6550 | ENTPD3 | 2 | Fluo4 | 0-5 min LG, 5-20 min HG, 20-25 min LG, 25 min-end KCl | 2.31721 |
| 6551 | ENTPD3 | 1 | Calbryte520 | 0-5 min LG, 5-20 min HG, 20-25 min LG, 25 min-end KCl | 0.37442 |
| 6551 | ENTPD3 | 2 | Calbryte520 | 0-5 min LG, 5-20 min HG, 20-25 min LG, 25 min-end KCl | 0.37585 |
| 6551 | ENTPD3 | 4 | Calbryte520 | 0-5 min LG, 5-20 min HG, 20-25 min LG, 25 min-end KCl | 0.37442 |
| 6551 | ENTPD3 | 5 | Calbryte520 | 0-5 min LG, 5-20 min HG, 20-25 min LG, 25 min-end KCl | 0.37728 |
| 6551 | ENTPD3 | 6 | Calbryte520 | 0-5 min LG, 5-20 min HG, 20-25 min LG, 25 min-end KCl | 0.37585 |
| 6552 | ENTPD3 | 1 | Calbryte520 | 0-5 min LG, 5-20 min HG, 20-27 min LG, 27 min-end KCl | 0.37585 |
| 6552 | ENTPD3 | 4 | Calbryte520 | 0-5 min LG, 5-20 min HG, 20-25 min LG, 25 min-end KCl | 0.37727 |
| 6552 | ENTPD3 | 5 | Calbryte520 | 0-5 min LG, 5-20 min HG, 20-25 min LG, 25 min-end KCl | 0.37585 |
| 6558 | ENTPD3 | 3 | Calbryte520 | 0-5 min LG, 5-20 min HG, 20-30 min LG, 30 min-end KCl | 0.37727 |
| 6558 | ENTPD3 | 4 | Calbryte520 | 0-5 min LG, 5-20 min HG, 20-30 min LG, 30 min-end KCl | 0.37585 |
| 6562 | RL | 2 | Calbryte520 | 0-5 min LG, 5-20 min HG, 20-30 min LG, 30 min-end KCl I | 4.38736 |
| 6562 | ENTPD3 | 3 | Calbryte520 | 0-5 min LG, 5-20 min HG, 20-30 min LG, 30 min-end KCl | 0.37443 |
| 6562 | ENTPD3 | 4 | Calbryte520 | 0-5 min LG, 5-20 min HG, 20-30 min LG, 30 min-end KCl | 0.37443 |
| 6563 | ENTPD3 | 2 | Calbryte520 | 0-5 min LG, 5-20 min HG, 20-30 min LG, 30 min-end KCl | 0.37585 |
| CV14 | ENTPD3 | 6 | Calbryte520 | 0-6 min LG, 6-21 min HG, 21-31 min LG, 31 min-end KCl | 0.37585 |
| CV14 | ENTPD3 | 7 | Calbryte520 | 0-5 min LG, 5-20 min HG, 20-30 min LG, 30 min-end KCl | 0.37727 |
| 6569 | ENTPD3 | 5 | Calbryte520 | 0-5 min LG, 5-20 min HG, 20-30 min LG, 30 min-end KCl | 0.37585 |
| 6569 | ENTPD3 | 6 | Calbryte520 | 0-5 min 30s LG, 5 min 30s-20 min 30s HG, 20 min 30s-30 min 30s LG, 30 min 30s-end KCl | 0.37585 |
| CV15 | ENTPD3 | 1 | Calbryte520 | 0-5 min LG, 5-20 min HG, 20-30 min LG, 30 min-end KCl | 0.37585 |
| CV15 | ENTPD3 | 2 | Calbryte520 | 0-5 min LG, 5-20 min HG, 20-30 min LG, 30 min-end KCl | 0.37585 |
| CV15 | ENTPD3 | 5 | Calbryte520 | 0-5 min LG, 5-20 min HG, 20-30 min LG, 30 min-end KCl | 0.37585 |
| CV15 | ENTPD3 | 6 | Calbryte520 | 0-5 min LG, 5-20 min HG, 20-30 min LG, 30 min-end KCl | 0.37585 |
| 6573 | ENTPD3 | 3 | Calbryte520 | 0-5 min LG, 5-20 min HG, 20-30 min LG, 30 min-end KCl | 0.37585 |
| 6573 | ENTPD3 | 4 | Calbryte520 | 0-5 min LG, 5-20 min HG, 20-30 min LG, 30 min-end KCl | 0.37585 |
| 6578 | ENTPD3 | 2 | Calbryte520 | 0-5 min LG, 5-20 min HG, 20-30 min LG, 30 min-end KCl | 0.37585 |
| 6578 | ENTPD3 | 3 | Calbryte520 | 0-5 min LG, 5-20 min HG, 20-30 min LG, 30 min-end KCl | 0.37585 |
| 6578 | ENTPD3 | 4 | Calbryte520 | 0-5 min LG, 5-20 min HG, 20-30 min LG, 30 min-end KCl | 0.37585 |
| 6579 | ENTPD3 | 1 | Calbryte520 | 0-5 min LG, 5-20 min HG, 20-30 min LG, 30 min-end KCl | 0.37727 |
| 6579 | ENTPD3 | 4 | Calbryte520 | 0-5 min LG, 5-20 min HG, 20-30 min LG, 30 min-end KCl | 0.37727 |
| 6579 | ENTPD3 | 7 | Calbryte520 | 0-5 min LG, 5-20 min HG, 20-30 min LG, 30 min-end KCl | 0.39293 |
| 6582 | ENTPD3 | 1 | Calbryte520 | 0-5 min LG, 5-20 min HG, 20-30 min LG, 30 min-end KCl | 0.37585 |
| 6583 | ENTPD3 | 1 | Calbryte520 | 0-5 min LG, 5-20 min HG, 20-30 min LG, 30 min-end KCl | 0.37727 |
| 6583 | ENTPD3 | 3 | Calbryte520 | 0-5 min LG, 5-20 min HG, 20-30 min LG, 30 min-end KCl | 0.37727 |
| 6583 | ENTPD3 | 4 | Calbryte520 | 0-5 min LG, 5-20 min HG, 20-30 min LG, 30 min-end KCl | 0.37585 |
| 6584 | ENTPD3 | 7 | Calbryte520 | 0-5 min LG, 5-20 min HG, 20-30 min LG, 30 min-end KCl | 0.37727 |
| 6584 | ENTPD3 | 9 | Calbryte520 | 0-5 min LG, 5-20 min HG, 20-30 min LG, 30 min-end KCl | 0.37727 |
| 6586 | ENTPD3 | 1 | Calbryte520 | 0-5 min LG, 5-20 min HG, 20-30 min LG, 30 min-end KCl | 0.37442 |
| 6586 | ENTPD3 | 2 | Calbryte520 | 0-5 min LG, 5-20 min HG, 20-30 min LG, 30 min-end KCl | 0.37728 |
| 6586 | ENTPD3 | 3 | Calbryte520 | 0-6 min LG, 6-21 min HG, 21-31 min LG, 31 min-end KCl | 0.37728 |
| 6588 | ENTPD3 | 1 | Calbryte520 | 0-5 min LG, 5-20 min HG, 20-30 min LG, 30 min-end KCl | 0.37727 |
| 6588 | ENTPD3 | 2 | Calbryte520 | 0-5 min LG, 5-20 min HG, 20-30 min LG, 30 min-end KCl | 0.37727 |
| 6588 | ENTPD3 | 3 | Calbryte520 | 0-5 min LG, 5-20 min HG, 20 min-30 min 40s LG, 30 min 40s-end KCl | 0.37727 |
| 6588 | ENTPD3 | 4 | Calbryte520 | 0-5 min LG, 5-20 min 30s HG, 20 min 30s-30min 30s LG, 30 min 30s-end KCl | 0.37727 |
| 6590 | ENTPD3 | 2 | Calbryte520 | 0-5 min LG, 5-20 min HG, 20-30 min LG, 30 min-end KCl | 0.37585 |
| 6590 | ENTPD3 | 3 | Calbryte520 | 0-5 min LG, 5-20 min HG, 20-30 min LG, 30 min-end KCl | 0.37585 |
| 6590 | ENTPD3 | 5 | Calbryte520 | 0-5 min LG, 5-20 min HG, 20-30 min LG, 30 min-end KCl | 0.37727 |
| 6591 | ENTPD3 | 6 | Calbryte520 | 0-5 min LG, 5-20 min HG, 20-30 min LG, 30 min-end KCl | 0.37727 |
| 6591 | ENTPD3 | 9 | Calbryte520 | 0-5 min LG, 5-20 min HG, 20-30 min LG, 30 min-end KCl | 0.37585 |
| 6591 | ENTPD3 | 10 | Calbryte520 | 0-5 min LG, 5-20 min HG, 20-30 min LG, 30 min-end KCl | 0.37585 |
| 6597 | ENTPD3 | 1 | Calbryte520 | 0-5 min LG, 5-20 min HG, 20-30 min LG, 30 min-end KCl | 0.37727 |
| 6597 | ENTPD3 | 3 | Calbryte520 | 0-5 min LG, 5-20 min HG, 20-30 min LG, 30 min-end KCl | 0.37728 |
| 6598 | ENTPD3 | 1 | Calbryte520 | 0-5 min LG, 5-20 min HG, 20-30 min LG, 30 min-end KCl | 0.37727 |
| 6598 | ENTPD3 | 2 | Calbryte520 | 0-5 min LG, 5-20 min HG, 20-30 min LG, 30 min-end KCl | 0.37585 |
| 6598 | ENTPD3 | 3 | Calbryte520 | 0-5 min LG, 5-20 min HG, 20-30 min LG, 30 min-end KCl | 0.37585 |
| 6598 | ENTPD3 | 4 | Calbryte520 | 0-5 min LG, 5-20 min HG, 20-30 min LG, 30 min-end KCl | 0.37727 |

**Supplementary Table 3:**
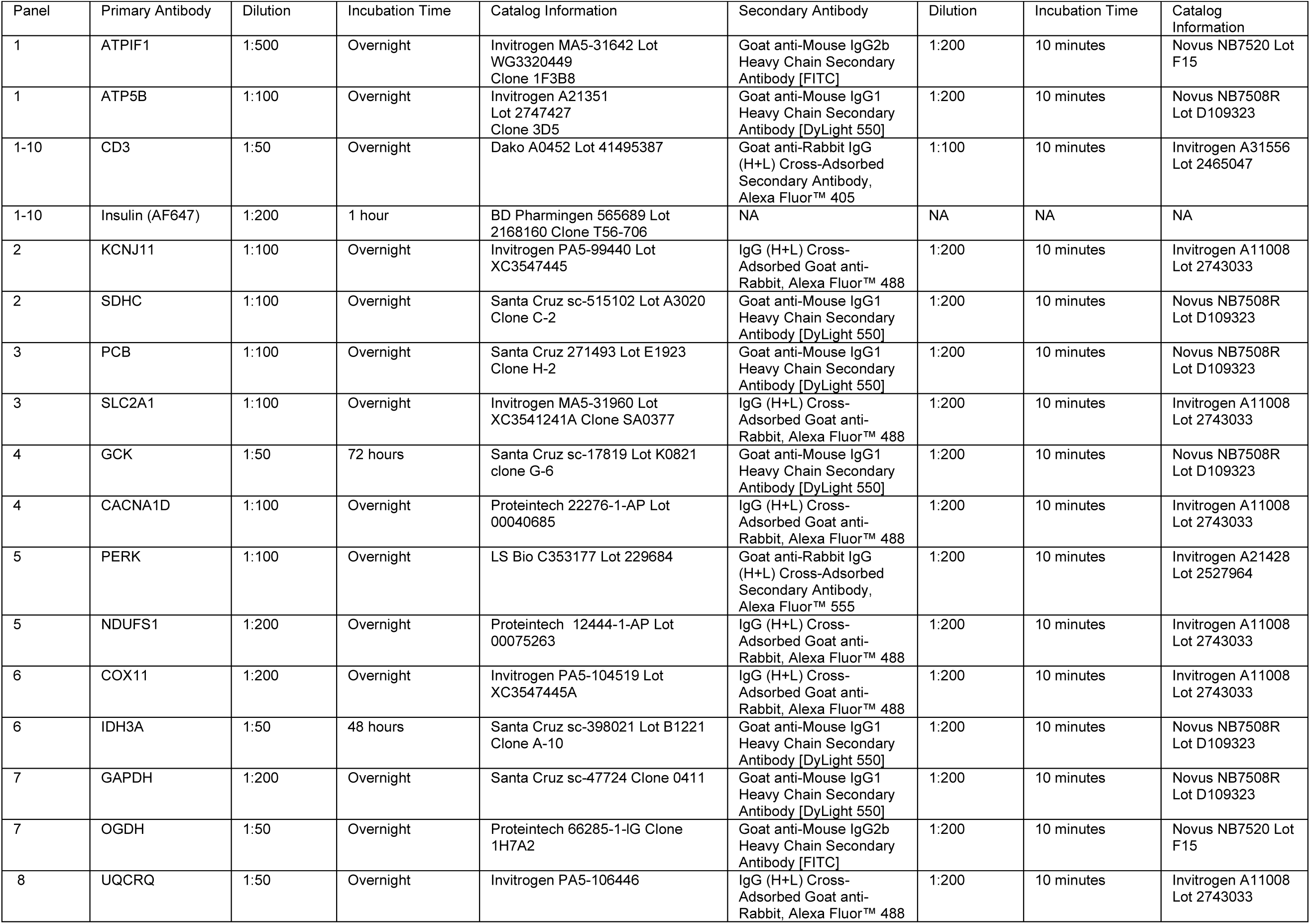
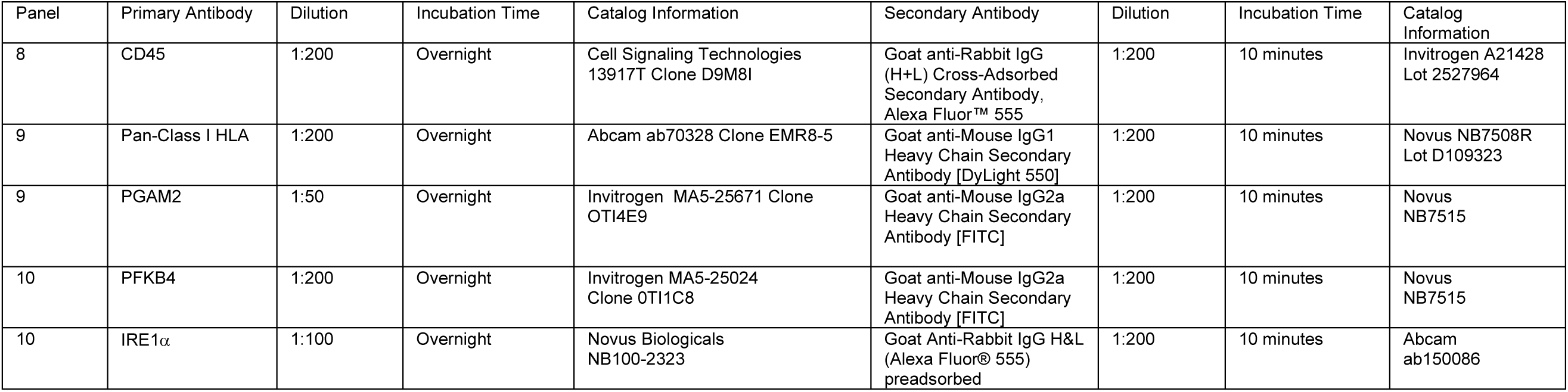
FFPE staining panels. Staining panels and protocols used for the immunofluorescent staining of FFPE pancreas tissue sections.

